# DIANNeR: label-free RIME resolves lineage-restricted Glucocorticoid Receptor interactomes in normal human tissues

**DOI:** 10.1101/2025.08.07.669166

**Authors:** Weiye Zhao, Thomas F Grimes, Susanna F Rose, Jack Stenning, Chris Taylor, Chloë Baldreki, Iain Goulding, Ros Duke, Simon C Baker, Marcela Montes de Oca, Aparna D Sinha, Jenny Hinley, James M Fox, Paul M Kaye, Jennifer J Gomm, Louise J Jones, Elisabetta Marangoni, Bruno M Simões, Robert B Clarke, Jennifer Southgate, Katherine S Bridge, Adam Dowle, Andrew N Holding

## Abstract

Understanding how transcription factors execute tissue-specific programmes requires defining their protein interaction networks in physiologically relevant contexts. However, profiling normal untransformed cells presents fundamental challenges due to intrinsically limited input material. To meet this challenge we integrated data-independent acquisition (DIA), ion mobility separation, and library-free analysis to achieve a 2-fold increase in detection without loss in enrichment.

Applied to the glucocorticoid receptor (GR), a ubiquitously expressed nuclear receptor driving pleiotropic responses to standard-of-care anti-inflammatory therapeutics, DIANNeR (DIA-NN enabled RIME) resolves distinct, context-dependent networks across breast, ureter and blood in normal and transformed contexts. Key findings include: detection of a HOXA5–GR interaction in normal epithelium which was undetected in malignant models; an epithelial-restricted SMARCD3–GR interaction with prognostic relevance; and a FOXP3–BCL11B–GR network in primary CD4^+^ T cell populations undetected by data dependant acquisition (DDA). By defining normal GR tissue interactomes, DIANNeR provides the essential comparator for interpreting network rewiring that drives disease.

## Main

Transcription factor (TF) activity depends not only on the protein itself, but also on the surrounding context-specific interaction networks. Despite this, our understanding of human TF interactomes remains heavily skewed towards immortalised cell lines and cancer models,presenting a critical problem: without a normal human baseline, how are pathological changes in disease states actually defined? Human tissue is predominantly sourced through finite donations of surplus material from necessary medical procedures. These non-renewable samples preclude the iterative optimisation and scale-up typically required for mass spectrometry-based proteomic interaction studies (interactomics). Consequently, the research field requires a validated, high-sensitivity framework capable of delivering comprehensive data from low amounts of donor material.

While Rapid Immunoprecipitation Mass spectrometry of Endogenous proteins (RIME)^1^ and its quantitative derivatives^2^ have established chromatin-bound interactome profiling of transcription factors as a tractable approach in immortalised cell lines and tumourigenic material^3^, they have yet to resolve the normal tissue bottleneck. Generating high-quality RIME data from normal tissues, which are non-proliferative and available only in limited quantities, has thus remained intractable compared to immortalised models.

Given the practical and ethical constraints on finite human tissues, our objective was to establish the analytical framework capable of resolving TF interactomes from patient material.

Here we present, DIANNeR (DIA-NN enabled RIME *or* Data-Independent Acquisition by Neural Networks–enabled Rapid Immunoprecipitation Mass Spectrometry of Endogenous Proteins, **Figure 1A**), a framework specifically designed for acquisitions of chromatin-bound complexes from scarce, untransformed cellular material. DIANNeR achieves this objective by leveraging data-independent acquisition (DIA) mass spectrometry combined with trapped ion mobility separation^4^ to overcome the stochastic sampling bottleneck inherent in DDA. These technologies are supported by pairing library-free spectral analysis via DIA-NN with the FragPipe-Analyst pipeline^5^, eliminating the need for additional sample material for spectral library generation while ensuring a generalisable output. Benchmarking in primary CD4^+^ T cells shows that DIANNeR approximately doubles the recovery of significantly enriched GR interacting proteins over DDA.

**Figure 1.**
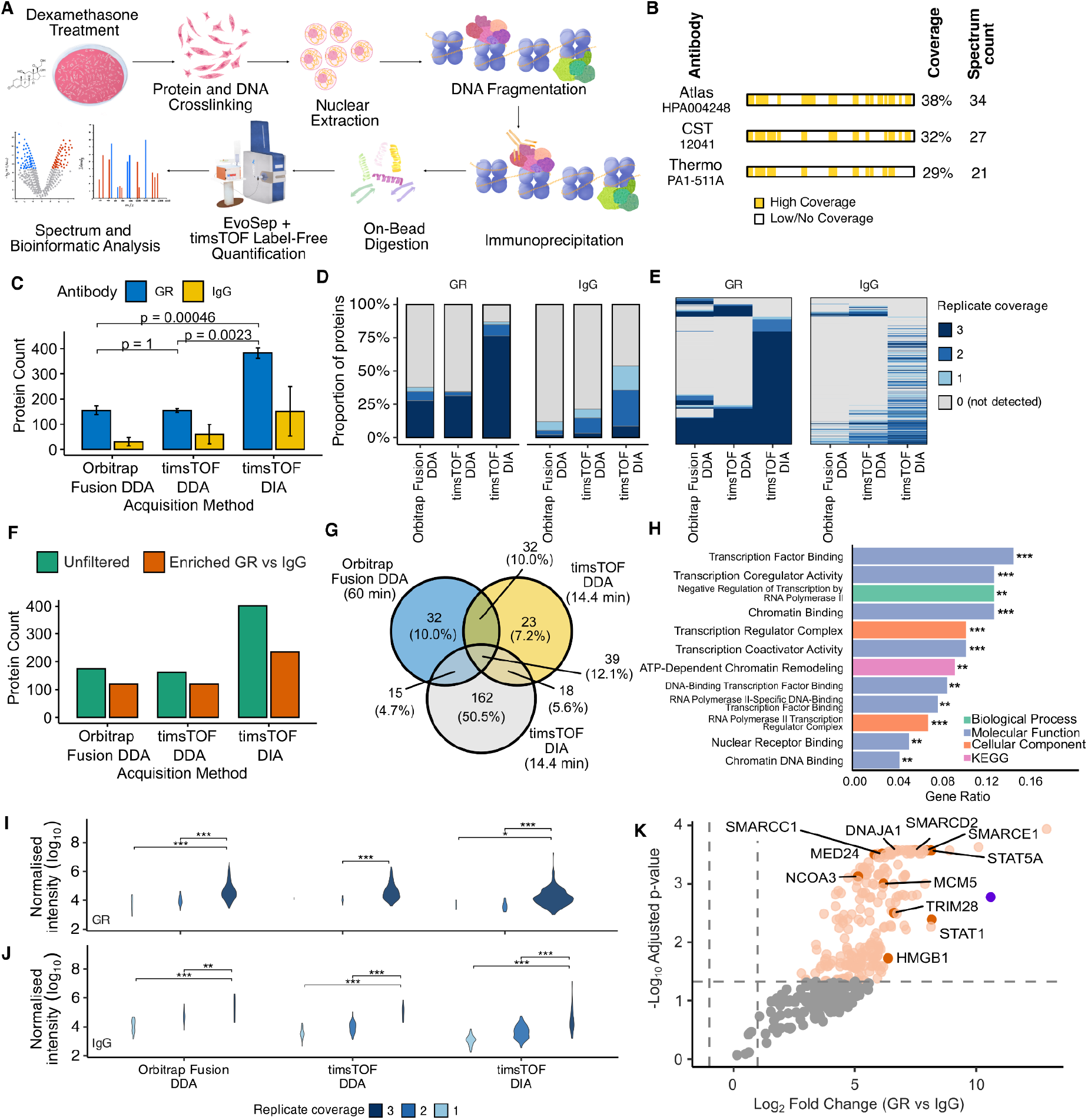
DIANNeR provides high-depth, reproducible TF interactomes from primary tissues. **(A)** Schematic overview of the DIANNeR workflow. **(B)** Antibody benchmarking: peptide spectrum count and sequence coverage on Glucocorticoid Receptor (GR) resulting from DDA-MS analysis of candidate GR antibodies. **(C)** Total protein identifications in GR RIME samples by acquisition method. DIANNeR identified significantly more proteins than either DDA approach (*t*-test, *n* = 3). **(D)** Data completeness: proportion of proteins detected across 1, 2, or 3 replicates for GR and IgG pull-downs. **(E)** Heatmap of replicate detection patterns for GR- and IgG-associated proteins. Colour intensity indicates the number of replicates where the protein was quantified. **(F)** Comparison of the number of proteins identified before and after IgG negative-control filtering demonstrates the increase in detected GR interactors is maintained after removal of background contaminants (LFC > 2, adjust-P < 0.05). **(G)** Venn diagram illustrating the overlap in proteomic coverage of GR interactomes across the three acquisition methods (LFC > 2, adjust-P < 0.05). **(H)** Enriched nuclear receptor, chromatin and transcription GO and KEGG terms from analysis of GR interactors in CD4^+^ T cells detected by DIA-RIME. **(I)** Relationship between protein abundance and replicate detection in GR pull-down samples. Log_10_-transformed intensity values were median normalised using a shared core proteome. Violin plots show the distribution of normalised intensities for proteins detected in 1, 2, or 3 replicates. Maximum width is proportional to protein count. Statistical significance was determined by one-way ANOVA with Tukey HSD post-hoc testing; all comparisons are made against the N=3 group (* *p* < 0.05, ** *p* < 0.01, *** *p* < 0.001). **(J)** Protein abundance distributions stratified by replicate detection for IgG pull down samples. **(K)** Volcano plot showing protein enrichment in GR-RIME samples relative to IgG controls using DIANNeR. GR is shown in purple.

We demonstrate this framework using the glucocorticoid receptor (GR or NR3C1), a ubiquitous transcription factor that drives radically divergent, context-dependent responses, from inducing apoptosis during development and maturation of T lymphocytes^6^ to suppressing involution in mammary epithelium^7,8^. In the bladder, glucocorticoids regulate diverse epithelial processes, including the coordination of the nocturnal micturition pattern^9^ and epithelial cell plasticity^10^. As a primary regulator of human homeostasis, and the target of some of the lowest cost and most effective clinical interventions, the GR represents a highly relevant model for demonstrating DIANNeR’s ability to resolve lineage-restricted signalling that is lost or modified in immortalised cell lines. Mapping the GR interactome across three distinct human lineages –normal breast epithelial cells, normal human urothelial cells, and primary CD4^+^ T lymphocytes– identified distinct changes in the GR interaction network when contrasted to cancer cell lines and patient-derived xenografts. Notably, the non-tumorigenic spontaneously immortalised MCF10A cell line, frequently used as a model for normal tissue, reveals a distinct interactome from primary breast epithelial cells, highlighting the necessity of untransformed normal cells to capture native signalling. DIANNeR further resolved lineage-restricted associations previously invisible in cell-line models, including a HOXA5–GR interaction lost upon malignant transformation in breast tissue, an epithelial-specific SMARCD3–GR association, and a Treg-associated FOXP3–BCL11B–GR network in primary CD4^+^ T cells that was absent in Jurkat models.

## Results

### DIA-NN-enabled RIME (DIANNeR) overcomes stochastic sampling limits to map primary tissue interactomes

To establish an optimised workflow for normal tissue analysis, we leveraged the trapped ion mobility separation (TIMS) of the Bruker timsTOF platform within a DIA workflow. TIMS provides an additional dimension of separation (Collisional Cross Section, CCS) to deconvolute the complex backgrounds of primary human samples. Additionally, after benchmarking three GR antibody candidates, HPA004248 was selected for superior bait and interactome recovery (**Figure 1B**, Online Methods).

For benchmarking, we performed a comparative GR DIA- and DDA-RIME assay using normal isolated human CD4^+^ T lymphocytes from three separate individuals and analysed all samples on each platform to prevent bias from varying pull-down efficiency. Our investigation, which involved parallel label-free quantitative analysis on three acquisition platforms (Fusion-DDA, timsTOF-DDA and timsTOF-DIA), enabled direct comparison to established DDA-RIME on the Orbitrap Fusion and comparison of DDA and DIA strategies on the timsTOF platform. DDA and DIA acquisition on the timsTOF followed identical chromatographic separation running a matched EvoSep 100 SPD LC gradient applied to identical physical samples split from each biological RIME replicate to ensure a paired, platform-matched comparison. The Orbitrap acquisition was aligned with established LC and DDA protocols. To our knowledge, this represents the first quantitative comparison of DIA-RIME (incorporating TIMS) against DDA workflows in primary human cells.

To isolate target-specific interactors, normal rabbit IgG was employed as an isotype control alongside GR immunoprecipitations, accounting for tissue-specific background in the IP-MS workflow^11^. Comparison of quantified protein level data demonstrated the DIA acquisition mode identified a significantly greater number of proteins than the DDA mode (t-test, n = 3, *p* < 0.05), using a minimum of two peptides per protein and a 1% FDR threshold (**Figure 1C**).

To ensure that this expanded interactome was driven by consistent, reproducible protein detection rather than expanding low-confidence background signal, we compared replicate consistency across the acquisition strategies. Our DIA timsTOF workflow showed a substantial increase in the proportion of proteins detected in all three replicates relative to both DDA Orbitrap Fusion and DDA timsTOF approaches, for both GR pulldowns and IgG controls **(Figure 1D)**. An analysis of per-protein detection patterns revealed that this increase was not the result of rescuing missing values, but from the expansion of the detectable interactome **(Figure 1E)**.

The enhanced proteome coverage via DIA acquisition was further supported by downstream analysis using FragPipe-Analyst^5^, which identified a 2-fold increase in the number of proteins significantly enriched over the IgG control in DIA datasets when compared to either of the DDA acquisitions (**Figure 1F**). GR itself was among the most abundant and significantly enriched proteins in the DIA-RIME dataset. Comparative analysis of significant GR interactions across the three platforms revealed that over 12% of proteins were commonly identified by all methods, with over 80% of all detected proteins recovered using our DIA based protocol **(Figure 1G)**. To quantify the biological quality of these expanded interactomes, we inferred specificity from curated-interactor databases, replicate completeness, and IgG controls. Performing hypergeometric testing against the BioGRID database we determined the enrichment of known GR interactors within each dataset^12^. Both DDA acquisition platforms demonstrated enrichment of known interactors (*p* = 0.04 and 0.013), while the DIANNeR-recovered interactome exhibited substantial improvement in statistical confidence (*p* = 1.36 × 10^-5^). These results support that the additional DIA-detected proteins are consistent with biologically validated components of the GR complex. Functional enrichment analysis of DIA-specific proteins further aligned with known GR biology, supporting the biological relevance of the detected interactome **(Figure 1H)**.

We next assessed the relationship between replicate detection and protein abundance in the GR DIANNeR samples. In the majority of datasets, proteins detected in fewer replicates were associated with a significantly lower mean intensity, consistent with intensity-dependent missingness. Importantly, the distribution of protein intensities from our DIA analysis demonstrated an increased detection across the entire intensity range, indicating that these detections are not solely attributable to stochastic sampling near the detection limit. **(Figure 1I)**. Analysis of the IgG samples **(Figure 1J)**, showed a correlation between missingness and lower abundance; however, this was associated with a high number of low abundance proteins in line with expectations of non-specific control pulldown.

Critically, the increased detection depth of DIANNeR translated to the recovery of several unique GR-interactors identified within BioGRID, including NCOA3, HDAC2 and SMARCC1. Notably, these coregulators were detected with high abundance in the DIA dataset **(Figure 1K)**, yet remained absent across both DDA acquisition approaches. As these benchmarks utilised identical physical samples of each biological replicate, this divergence cannot be attributed to biological variation.

Together, these results demonstrate that DIANNeR provides superior data completeness over DDA-RIME strategies, and resolves high-abundance coregulators that are otherwise lost to stochastic sampling.

### Clustering GR interactome by protein intensity of DIANNeR samples demonstrates tissue-specific differences

Having established the superior sensitivity of DIANNeR, we next evaluated its ability to resolve context-specific regulatory complexes across a panel of three primary human tissues and matched cancer cell lines **(Figure 2A)**. After consolidating two independent acquisition batches using pooled controls to monitor batch effects, we identified a total of 7914 proteins across our panel. The resulting protein-centric dataset is hereafter referred to as the DIANNeR GR dataset. Principal component analysis (PCA) revealed samples clustered primarily by pull-down specificity (IgG vs GR-specific antibody) across PC1 and PC2 **(Figure 2B)**. GR-specific immunoprecipitations were separated from IgG controls for all cellular models, confirming the specificity of GR interactome enrichment. Additionally, pooled samples from the second batch of analysis clustered with their original samples, indicating minimal batch effects. Visualisation of PC3 through to PC8 systematically isolated individual cell types and tissues or origin **(Extended Data Figure 1A-C)**.

**Figure 2.**
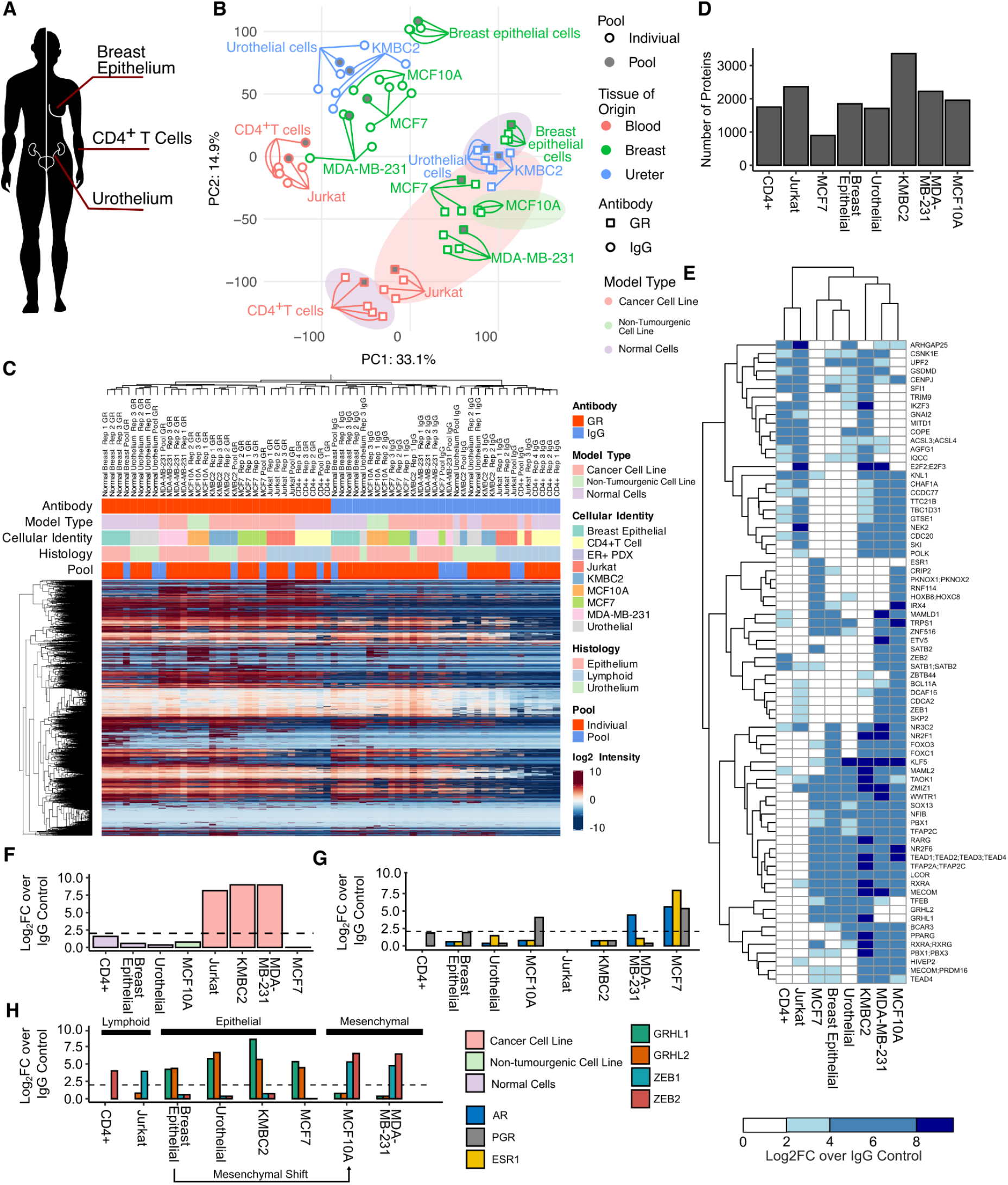
Clustering of samples on detected protein intensity demonstrates reproducibility of DIA-NN enabled RIME (DIANNeR) for GR interactome analysis and LFC analysis identifies context specific interactions. **(A)** Sites of primary tissue collection. **(B)** PCA analysis of protein intensities across all samples demonstrates clustering by antibody and tissue of origin for all cell-based IPs. **(C)** Hierarchical clustering (Euclidean distance) of protein intensities for all samples from both batches clustered. GR RIME Pooled samples included in the second batch reproducibly clustered with their corresponding individual samples from the first batch. For GR specific pulldowns, clustering by epithelial vs lymphoid cell types dominates. The strength of IgG RIME samples clustering was primarily defined by tissue of origin rather than interactome. **(D)** The number of unique proteins significantly enriched over IgG control for each model (LFC > 2 and *p*-adjust < 0.01). **(E)** Heatmap of lineage-defining transcription factors driving sample clustering. DIANNeR resolves distinct sets of coregulators that distinguish primary tissues from their immortalised counterparts. **(F)** Enrichment of E2F2/E2F3 across the panel; interaction with GR is uniquely identified in proliferative cancer models but absent in healthy primary tissues. **(G)** Enrichment of steroid receptors (AR, PGR, ESR1) matches established tissue biology, validating the framework’s ability to capture known hormonal signalling nodes. **(H)** DIANNeR identifies an epithelial-to-mesenchymal transition in the GR interactome of the normal MCF10A cell line, characterised by the loss of GRHL1/2 and gain of ZEB1/2 interactions compared to primary breast epithelium.

GR interactomes for lymphoid samples formed a distinct clustering separate from epithelial-derived samples. Similarly, the GR interactome from urothelium most closely resembled that of the bladder cancer cell line KMBC2. Conversely, the non-tumourigenic breast model, MCF10A, failed to cluster with primary breast epithelium, instead associating with the interactomes of the breast cancer cell lines (**Figure 2B**).

To complement our PCA, we undertook hierarchical clustering of samples by protein intensity. This analysis independently confirmed the accuracy and reproducibility of the method, with pooled samples again clustering with their corresponding initial batches **(Figure 2C)**. Consistent with our PCA results, Jurkat and normal CD4^+^ T cells clustered by tissue of origin for the GR specific pulldown, while the GR interactome for normal breast epithelium and urothelium is clustered together distinct from established breast and bladder cancer cell lines. Most notably, the GR interactome of the non-tumourigenic breast cell line MCF10A clustered most closely with the TNBC cell line MDA-MB-231 rather than primary epithelium, in clear preference over the ER+ breast cancer model MCF7 (**Figure 2C**). Linear modelling confirmed that inherent protocol differences between suspension and adherent models had limited influence on the overall profiles, supporting that the observed separations are driven by cell biology **(Extended Data Figure 1D)**.

### Differential analysis of GR interactomes across normal cells and cancer cell lines highlights tissue-specific GR interactions and functions

The differential selection and abundance of coregulators within the GR interactome across tissues is key in shaping the unique, context dependent responses to glucocorticoids ^13^. To quantitatively map these tissue-specific signatures, we performed differential protein abundance analysis using the FragPipe-Analyst pipeline. Fold enrichment of GR-interacting proteins was calculated for each protein in the GR IP group relative to the IgG control group using the linear model implemented in FragPipe-Analyst **(Results File S1)**. We established the high-confidence GR interactors for each model (LFC > 2, and *p-*adjust < 0.01, **Figure 2D)** and prioritised the most informative features using PCA to establish the top contributing interactors across PCA1-3 **(Extended Data Figure 2, Figure 2E).**

The prioritised GR interactome heatmap **(Figure 2E)** showed distinct clustering patterns underlying the biology of the models. The two lymphoid models clustered separately from the other tissues, while ER+ MCF7 and normal cells from the two epithelial tissues formed a second cluster. Notably, the non-tumourigenic but immortalised MCF10A line again failed to cluster with normal breast epithelium, instead associating with TNBC model MDA-MB-231. Overall, we believe that the prioritised GR interactors outlined in **Figure 2E** provide an informative overview of the model-specific GR interactomes.

#### The GR interactome distinguishes proliferative cancer models via E2F recruitment

Differential binding analysis identified a high-affinity recruitment of the E2F2/E2F3 protein group (LFC > 8) specifically within the Jurkat, KMBC2, and MDA-MB-231 cancer cell lines **(Figure 2F)**. This interaction was conspicuously absent in all healthy primary tissues. While MCF7 is a transformed cancer line, its GR interactome also lacked this E2F signature, instead maintaining an interactome that aligns more closely with normal epithelial cells. This divergence suggests that the E2F-GR interaction is not a universal feature of transformation, but rather a specific marker for high-proliferative or aggressive cancer states. Jurkat cells are considered resistant to GR mediated-apoptosis^14,15^, and early (1-hour) treatment windows capture a phase where GR engages E2F transcription factors similarly to other highly proliferative cancer models. Jurkat cells are highly proliferative with a reported doubling time of 26 hours^16^, similar to that reported for MDA-MB-231 and KMBC^17,18^. Our findings demonstrate that DIANNeR can resolve these distinct pathological signatures, distinguishing the epithelial-like interactome of MCF7 from the more aggressive profiles of TNBC and lymphoid models.

#### GR interaction profiles track the loss of epithelial identity

To validate our findings against established steroid biology, we confirmed the MCF7-specific interaction between GR and ERα (ESR1), consistent with the known GR-ERα regulatory axis in ER+ breast cancer. **(Figure 2G)**^19–22^ Beyond this, we identified specific interactions with the androgen (AR) and progesterone (PGR) receptors that align with tissue context: AR was restricted to MCF7 and MDA-MB-231, while PGR recruitment was observed in both MCF10A and MCF7. We further observed that GR engages with transcription factors that define lineage identity and the transition toward a mesenchymal state. GRHL1 and GRHL2, TFs associated with epithelial identity and differentiation, interacted with GR in MCF7, normal breast epithelial and normal urothelial cells, and KMBC2 cells. By contrast, GR preferentially interacted with ZEB1 and ZEB2, key drivers of epithelial-to-mesenchymal transition (EMT)^23^, in MDA-MB-231 cells, consistent with their mesenchymal phenotype^24^ (**Figure 2H)**.

We further noted the non-tumourigenic MCF10A line also displayed this ZEB1/2-GR interaction profile, representing a departure from the GRHL-centric interactome observed in the normal breast epithelium. This shift in coregulator preference matches the PCA and hierarchical clustering **(Figure 2B, E)** where the MCF10A GR interactome associates with the aggressive MDA-MB-231 cell-line. Consequently, these data indicate that under established culture conditions for these cell types, the GR interactome serves as a reporter of the underlying differentiation and mesenchymal state, revealing that GR-signalling in the immortalised model MCF10A is more reflective of a transformed state than previously appreciated.

### Integrative analysis of differential GR interactomes identifies tissue-specific features including an ERα-associated interactome in breast cancer cell lines

To investigate context-specific differences in the GR interactome across our breast models, we performed differential enrichment analysis between normal breast epithelial cells, two breast cancer cell lines (MDA-MB-231 and MCF7), and the non-tumourigenic cell line model MCF10A.

Differential binding analysis of MDA-MB-231 vs MCF7 GR interactomes **(Figure 3A)** revealed enhanced association of ZEB1 and ZEB2 with GR in the mesenchymal MDA-MB-231 line, consistent with our earlier PCA findings **(Figure 2E)**. Equally, we identified GRHL2 as one of the most significantly enriched GR interactors in the ER+ MCF7 cells, alongside significant enrichment of GRHL1. The equivalent comparison of the GR interactome in MCF7 cells to normal breast epithelial cells showed a similar enrichment for ERα and related proteins including FOXA1, GATA3, and GREB1 **(Figure 3B)**.

**Figure 3.**
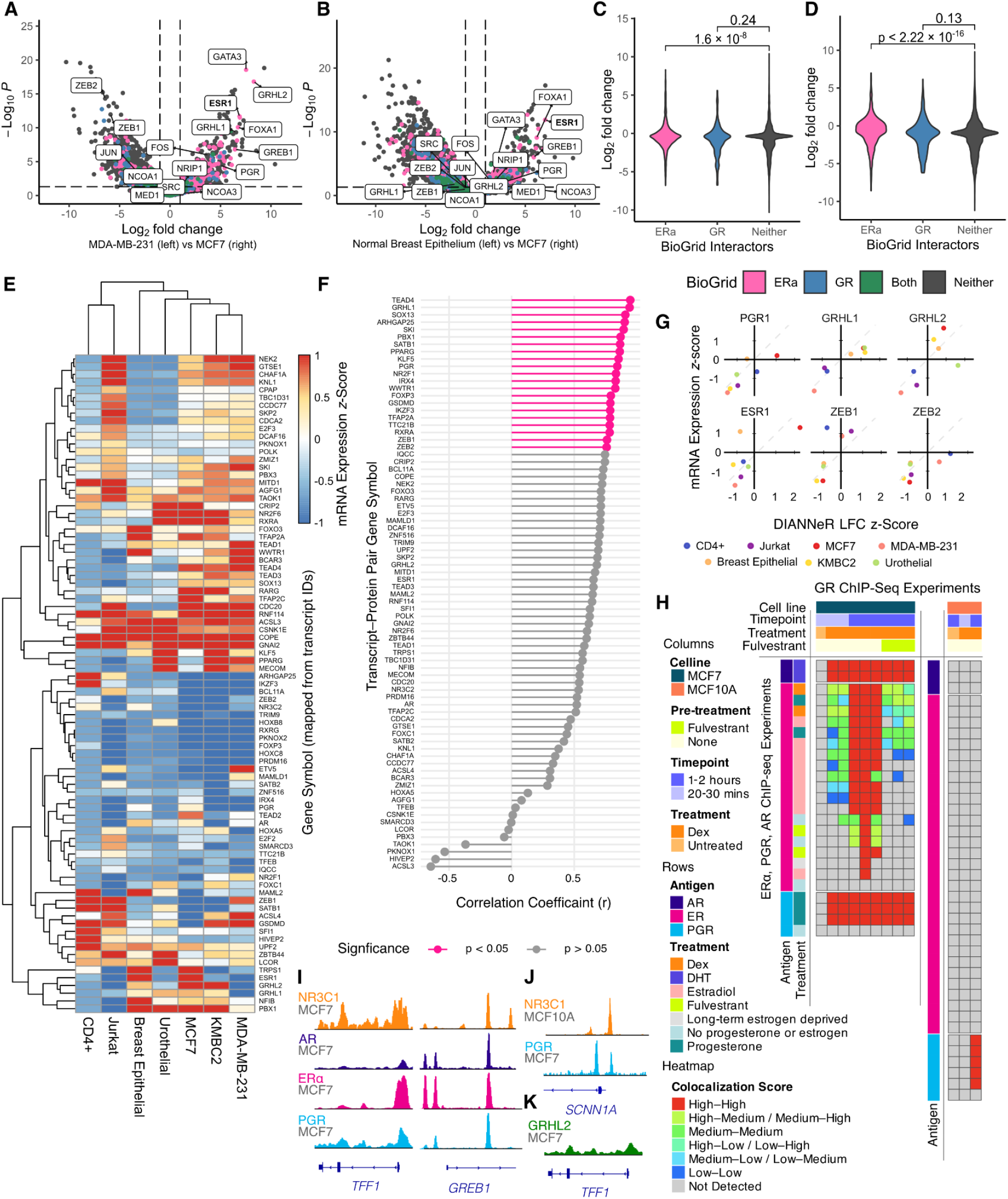
DIANNeR identifies specific differences in the GR interactome between breast cancer cell lines and normal breast epithelial cells. **(A)** Volcano plot comparing GR interactors in MCF7 and MDA-MB-231 cells, highlighting known ERα and GR-associated proteins. **(B)** Volcano plot comparison of the GR interactors in MCF7 cells and normal breast epithelium, highlighting known ERα and GR interacting proteins. **(C)** Violin plot for LFCs for GR interactors from comparison of MCF7 and MDA-MB-231 cell lines, stratified by their assignment as GR or ERα protein interactors as established from BioGrid. **(D)** Violin plot for LFCs for GR interactors from comparison of MCF7 and normal breast epithelium cells and their assignment as GR or ERα protein interactors as established from BioGrid. **(E)** z-score normalised transcript abundance for proteins most strongly associated with tissue-specific GR interactions. **(F)** Pearson correlation values for transcript-protein interaction pairs. Nominal correlation shown for p < 0.05; no correlation was significant after Bonferroni correction. **(G)** Scatter plots for individual transcript:protein–GR interaction z-score comparisons. The dotted line indicates the 45° diagonal where z-scores for expression and interaction values are equal. **(H)** (Left) Colocalisation heatmap quantifying the overlap between GR ChIP-seq peaks in MCF7 samples (columns) with AR, ERα and PGR (rows) and (Right) GR peaks from MCF10A samples (columns) compared against steroid receptor ChIP-seq datasets from MCF7 cells (columns). **(I)** Representative ChIP-seq tracks showing GR (GSM2154989), AR (GSM2797088), ERα (GSM1469980), PGR (GSM3417123) binding in MCF7 cells at the *TFF1* and *GREB1* loci. **(J)** Representative ChIP-seq tracks demonstrating colocalisation between GR binding in MCF10A cells (GSM2970418) and PGR binding in MCF7 cells at the SCNN1A locus. **(K)** Representative ChIP-seq track demonstrating GRHL2 (GSM2970418) at *TFF1* locus, consistent with steroid hormone receptor binding at the same loci. All genome-browser views correspond to the GRCh38 reference assembly.

Importantly, our DIANNeR pipeline recapitulated previous findings that GR and ERα interact in MCF7 cells ^20^. To assess this relationship, we stratified LFCs for GR- and ERα-interacting proteins as annotated in BioGRID. We observed a significant increase in LFCs of ERα-associated proteins in the MCF7 GR interactome relative to the MDA-MB-231 interactome, when compared to proteins not classified as GR or ERα interactors. No significant differences were observed in LFC between MCF7 and MDA-MB-231 for GR interacting proteins **(Figure 3C)**. We also detected a significant increase in LFCs of ERα-interacting proteins in MCF7 compared to normal breast epithelial cells **(Figure 3D)**.

### Bulk transcript levels do not uniformly predict the chromatin-bound GR interactome across models

To evaluate if transcriptional abundance predicted GR-protein associations, we integrated mRNA expression with DIANNeR-derived interacting profiles **(Figure 3E)**. No global correlation survived multiple testing corrections, indicating that bulk transcript levels do not uniformly dictate the GR interactome. However, acknowledging transcriptome-wide correction could mask genuine biological signals, we subsequently deployed a targeted candidate approach. Restricting our analysis of GR interactors to the core network identified by PCA **(Figure 2E)**, alongside *HOXA5, SMARCD3, FOXP3, PGR,* and *AR* based on the observations from our cell-type comparisons, revealed 21 proteins with nominal linear associations (r > 0.75, uncorrected *p* < 0.05) (**Figure 3F**). None of these associations retained significance following Benjamini–Hochberg adjustment.

The nominal relationships were most prominent among epithelial factors, including *TEAD4* (and its cofactor *WWTR1* ^25,26^), *PGR*, and *EMT* regulators *GRHL1* and *ZEB1/2*. Conversely, the association between *ESR1* transcript levels and the GR–ERα protein interaction did not meet nominal significance (**Figure 3G, Extended Data Figure 3A)**. Among lymphoid regulators, FOXP3 recruitment was coupled to its specific expression in CD4^+^ T-cell samples, whereas *IKZF3* and *SATB1* showed a broader lymphoid signature, with both expression and GR-association extending into Jurkat models **(Extended Data Figure 3B).**

Collectively, these findings demonstrate that transcript abundance is an unreliable predictor of physical GR recruitment. While baseline expression is a prerequisite for physical interaction, this decoupling is perhaps expected, given that cellular abundance alone does not govern nuclear receptor activity. Ultimately, the failure of these expression–interaction relationships to maintain significance under multiple-testing adjustment, even within a targeted approach, underscores that cofactor transcript expression cannot reliably predict functional GR association.

### DIANNeR interactions correspond to chromatin co-occupancy in breast models using public ChIP-seq datasets

To determine whether tissue-specific GR interaction profiles identified by DIANNeR correspond to colocalisation of transcription factors on the chromatin, we integrated our proteomic findings with GR, ERα, PGR and AR ChIP-seq datasets.

Analysis of ChIP-seq profiles for MCF7s cells revealed a high degree of colocalisation of GR, with all three nuclear receptors in the presence of dexamethasone (**Figure 3H**). Interestingly, recruitment of GR to sites typically bound by ERα was ERα-dependent. GR-ChIP-seq generated data in the presence of the selective estrogen receptor degrader (SERD) fulvestrant markedly reduced GR binding at sites bound by ERα in control conditions. In contrast, fulvestrant had a minimal effect on GR colocalisation at sites typically occupied by AR or PGR. The ChIP-seq coverage track demonstrating colocalisation of GR, ERα, PGR and AR is shown at *TFF1* and *GREB1* loci in **Figure 3I (Supplementary Figure S1).**

As MCF10A cells lack ERα and AR expression^27^, we assessed colocalisation between GR peaks in MCF10A with steroid receptor binding in MCF7. GR peaks in MCF10A displayed colocalisation with PGR binding sites from MCF7, but not with ERα or AR **(Figure 3J, Supplementary Figure S2)**. This finding aligns with our DIANNeR analysis, which detected PGR, but neither ERα nor AR, within the MCF10A GR interactome. Pairwise comparative analysis of the normal breast GR interactome reveals within-tissue rewiring on transformation and lineage-specific networks.

Beyond nuclear receptors, we confirmed colocalisation of GR binding with the GRHL2, TRPS1 and TEAD4 peaks in MCF7s (identified in **Figure 2E**). At the *TFF1* locus, GRHL2 demonstrated colocalised GR, at a site also bound by ERα, AR and PGR, and consistent with the established ERα–GRHL2 axis **(Figure 3K, Supplementary Figure S1A**)^28–31^. TRPS1 colocalisation required GR activation by dexamethasone, and GR binding at TRPS1 sites persisted in the context of fulvestrant **(Extended Data Figures 3C&D).** Together, these data validate the DIANNeR-predicted interactions and demonstrate that the GR cistrome is shaped by both stable and transient co-regulatory associations that are highly dependent on the cellular receptor context.

### Interactomic Profiling Across Breast Models Reveals the Rewiring of Normal Epithelial HOX-GR Networks Upon Transformation

To explore further changes in GR-associated networks across different breast models, we directly compared the GR interactome in MDA-MB-231 cells with that of normal breast epithelial cells. Consistent with our PCA results, GR in MDA-MB-231 cells showed stronger associations with the mesenchymal transcription factors ZEB1 and ZEB2, whereas normal breast epithelial cells exhibited enhanced interactions with the epithelial regulators GRHL1 and GRHL2. While we did not observe significant enrichment of ERα in normal cells over that in the MDA-MB-231s, likely reflecting the low abundance of ER-positive cells within breast reduction specimens, we did detect enrichment of GATA3, a pioneer factor for ERα and a key marker of breast epithelial identity ^32^ **(Figure 4A, left)**.

**Figure 4.**
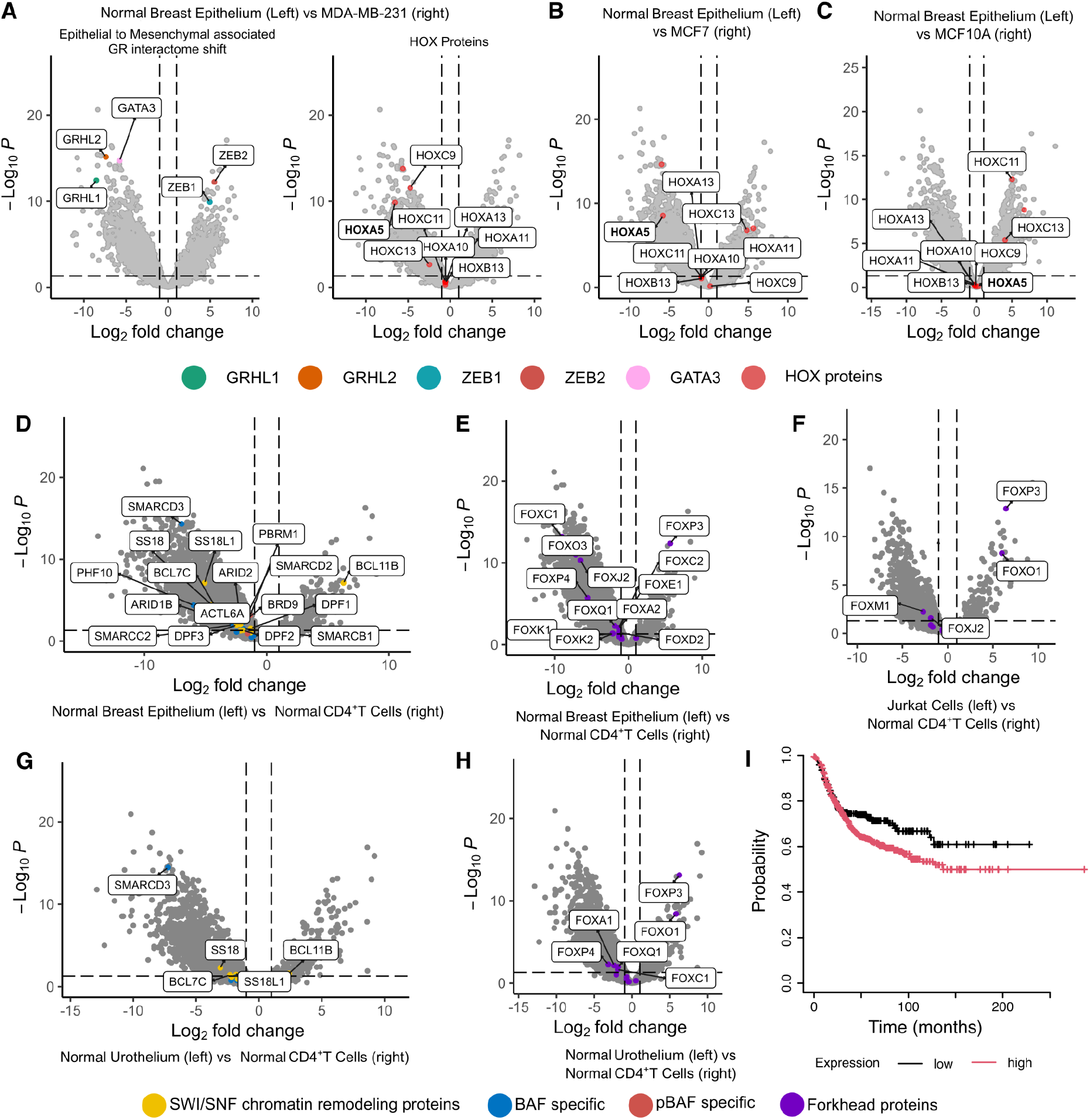
Comparison of lineage-specific GR interactomes reveals an epithelial-specific SMARCD3 axis and T_reg_ associated protein FOXP3 is enriched in healthy CD4+ T cell GR interactome but lost in Jurkat cells. **(A)** Volcano plot comparing GR interactors in MDA-MB-231 and normal breast epithelial cells, highlighting core lineage-associated proteins (*left)* and HOX protein interactions (*right*). **(B)** Volcano plot comparing GR interactors in MCF7 and normal breast epithelial cells, highlighting HOX protein interactions. **(C)** Volcano plot comparing GR interactors in MCF10A cells and normal breast epithelial cells, highlighting HOX protein interactions. **(D)** Volcano plot comparing GR interactors in normal breast epithelium and CD4+ T cells, highlighting proteins from the SWI/SNF complex with LFC > 2 and *p*-adjust < 0.05. **(E)** Volcano plot comparing GR interactors in normal urothelium and CD4+ T cells, highlighting SWI/SNF complex with LFC > 2 and *p*-adjust < 0.05. **(F)** Volcano plot comparing GR interactors in normal breast epithelium and CD4+ T cells, highlighting forkhead proteins. **(G)** Volcano plot comparing GR interactors in normal urothelium and CD4+ T cells, highlighting forkhead proteins. **(H)** Volcano plot comparing GR interactors in the Jurkat cell line and normal CD4+ T cells, highlighting forkhead proteins. **(I)** Kaplan-Meier survival analysis associates higher expression of *SMARCD3* with worse recurrence-free survival in ER-, HER2- patients (RFS; n=867, HR 1.32, p-value = 0.033).

In investigating differential GR interactions between normal tissues and cell line models, we identified tissue-dependent associations with HOX proteins. Notably, the protein HOXA5 was consistently enriched in the GR interactome of normal breast cells compared to both cancerous cell lines **(Figure 4A, right and 4B).** In contrast, no significant difference in HOXA5–GR interaction was observed when comparing normal cells with the non-tumourigenic MCF10A cell line **(Figure 4C)**; suggesting this interaction is preserved in normal and immortalised epithelium and diminished below detection in malignant models under our assay conditions.

### Cross-Lineage Baselines Resolve a Conserved Epithelial GR–SMARCD3 Axis and Treg-Specific GR-FOXP3 Networks within Normal CD4^+^ T Cell Populations

Given the divergent responses across different tissues, we next examined how SWI/SNF–GR interactions varied between epithelial and lymphoid contexts, reflecting the established role of mediating GR-driven transcriptional responses^33–35^. Within the CD4^+^ T cell lineage, BCL11B emerged as the most significantly enriched member of the SWI/SNF complex compared to breast epithelium **(Figure 4D).** Noting the reported interdependence of BCL11B and FOXP3 in regulatory T (Treg) cell function^36^, we explored GR interactions across the broader forkhead (FOX) protein family. Our targeted analysis revealed FOXP3 was the most significantly enriched Forkhead protein **(Figure 4F).** In contrast, comparison of our normal CD4^+^ T cells to the Jurkat T cell line **(Figure 4F)** revealed FOXP3 was absent from the GR network in the cell line.

To establish whether these differential interactomes reflected generalisable distinctions between lymphoid and epithelial lineages rather than breast-specific traits, we expanded our analysis to include normal urothelium. While the urothelium versus CD4^+^ T cell comparison identified fewer overall differential GR interactors than observed in the breast epithelium, we did find conserved and significant enrichment for the BAF-specific core component SMARCD3 **(Figure 4G)**. Equally, the normal lymphoid-specific enrichment of BCL11B and FOXP3 was maintained in both epithelial-versus-lymphoid comparisons **(Figure 4H)**, confirming that these distinct interaction profiles represent reproducible, lineage-specific networks.

To explore the clinical relevance of this lineage-restricted axis as an avenue for cell-type-specific therapeutic design, we examined *SMARCD3* expression across breast cancer subtypes. *SMARCD3* expression has previously been associated with improved prognosis in ER+ breast cancer ^37^, while our analysis of TNBC patients (ER −ve by IHC, HER2 −ve by array) revealed the opposite association; higher *SMARCD3* expression correlated with worse outcome (*p* = 0.033, n=867, **Figure 4I**) ^38,39^. These contrasting clinical trajectories reflect the known, subtype-specific outcomes of glucocorticoid signalling in breast cancer.

### Patient-derived tumour models support *in vivo* transcription factor network rewiring

To validate our cancer-associated findings in an *in vivo* setting, we undertook analysis of GR interactome from three independent ER+ breast cancer PDX models. We demonstrated enrichment of GR-interacting proteins over IgG control for each individual PDX by applying a threshold of LFC > 2, followed by overrepresentation analysis for relevant terms. All three PDX samples showed significant enrichment of terms related to nuclear receptor binding (GO:0016922), nuclear receptor-mediated steroid hormone signalling (GO:0030518), signalling by nuclear receptors (R-HSA-9006931) and transcription coregulator activity (GO:0003712) **(Figure 5A,B)**. Based on these results, we concluded that GR complexes were successfully enriched in all three models, thereby justifying their inclusion in subsequent combined analysis. Combining all three PDX models as biological replicates in FragPipe-Analyst, followed by filtering the combined samples on both LFC and significance (LFC > 2, *p*-adjust < 0.01), we observed pathway enrichment consistent with our individual PDX analyses **(Figure 5C)** which featured canonical GR interactors, including EP300, PRMT3, and NCOA1 **(Figure 5D)**.

**Figure 5.**
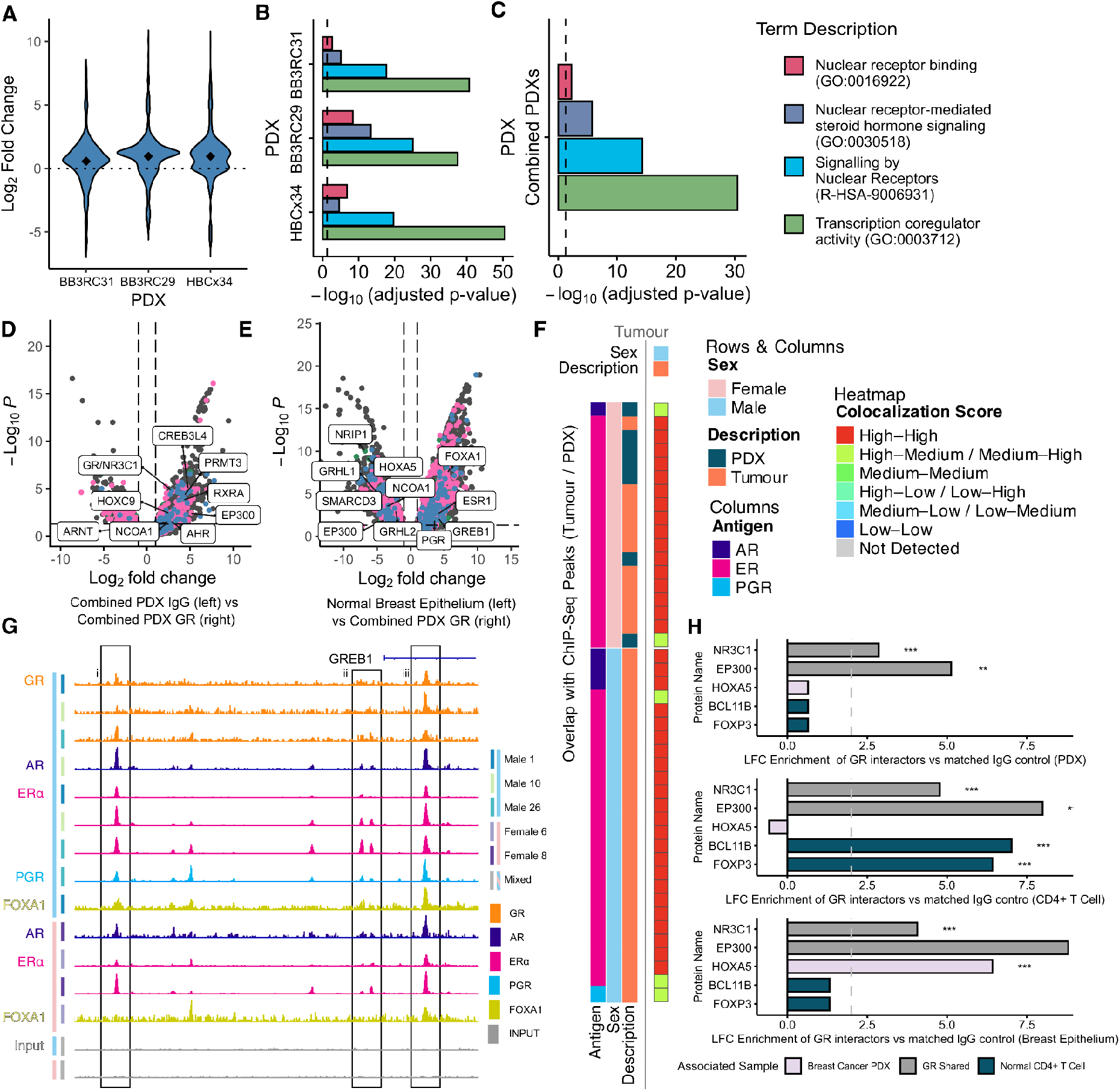
Characterization and tissue-specificity of the GR and ERα co-regulatory interactome in ER+ breast cancer PDX models. **(A)** Individual analysis of each PDX model revealed consistent enrichment of a subset of proteins in GR immunoprecipitates relative to IgG control. **(B)** Filtering proteins using LFC > 2 and *p*-adjust < 0.01 demonstrated significant enrichment for key terms related to the GR interactome for each PDX. **(C)** Filtering proteins using LFC > 2 and *p*-adjust < 0.01 demonstrated significant enrichment for key terms related to the GR interactome for combined PDX data, consistent with the findings for individual PDXs. **(D)** Volcano plot of the combined GR-IP data from all three PDX models, showing significant enrichment of GR (NR3C1) along with known GR-associated proteins including EP300, NCOA1 and RXRA. Known GR interactors are coloured blue, known ERα interactors are pink. **(E)** Volcano plot comparing GR interactomes from the PDX models and normal breast epithelial cells. Known GR and ERα interactors are coloured as in (C). **(F)** Colocalisation heatmap quantifying the overlap between GR ChIP-seq peaks in Patient Tumours (columns) with AR, ERα and PGR (rows) from tumour and PDX samples (columns). **(G)** Representative ChIP-seq tracks for GR, AR, ERa, PGR, and FOXA1 colocalisation across 5 patient tumours (GSE104399). **(H)** Comparison of LFC enrichment for selected lineage specific GR-associated proteins against matched IgG controls across combined PDX models, CD4+ T cells, and normal breast epithelium.

Direct comparison of the GR interactome from PDX models with that of normal breast epithelium recapitulated key findings from cell lines. Specifically, we identified significant enrichment of ERα-associated proteins including ERα (*p*-adjust = 0.0015), GREB1 *(p*-adjust = 0.0075) and FOXA1 (*p*-adjust = 3.6 × 10^-8^) in the PDX samples, over that of the normal epithelium. In contrast, HOXA5 was significantly enriched in normal epithelial samples (*p*-adjust = 5.5 × 10^-10^), consistent with our results comparing normal epithelium and cell lines **(Figure 5E)**. These data support a selective loss of the HOXA5–GR interaction during malignant transformation and the acquisition of an ERα-enriched GR complex in ER+ breast cancer.

Furthermore, we integrated these proteomic results with public ChIP-seq datasets^40^ from clinical and PDX ER+ breast tumour samples. Genomic GR binding consistently colocalised with ERα, PGR and AR peaks from ER+ tumours and PDX models irrespective of sex (**Figure 5F)**. Visualisation of individual ChIP-Seq tracks for 5 patient samples demonstrated GR colocalisation at *GREB1* locus **(Figure 5G)**, providing definitive *in vivo* genomic validation of our physical proteomic networks in a clinical context.

To definitively establish that DIANNeR resolves the physiological, context-dependent nature of transcription factor complexes *in vivo*, we mapped our lineage-specific topologies governing the GR interactome across normal tissue, and established cell lines to our PDX models. While significant enrichment of GR and its co-activator EP300 was highly conserved across all cellular contexts, the HOXA5–GR axis was enriched exclusively within the native normal breast epithelial context and lost within the malignant PDX models. Conversely, the BCL11B–FOXP3–GR lymphoid-specific axis was robustly and selectively maintained in primary CD4^+^ T cells, but was not significant within either the normal or malignant breast tissue settings **(Figure 5H).**

## Discussion

A central challenge in human cell biology is our reliance on immortalised lines as models. This limitation is particularly acute in breast tissue research, where human development and oncogenesis are poorly recapitulated by murine models. Further, the scarcity of normal primary tissues, combined with their restricted ability to expand *in vitro* without undergoing transformation severely limits routine optimisation. DIANNeR has demonstrated the ability to acquire biologically relevant chromatin-bound GR-interacting coregulators from untransformed cellular material. We demonstrate its broad generalisability by successfully mapping these interactomes across normal breast epithelium, urothelium, primary across normal breast epithelium, urothelium, primary CD4^+^ T lymphocytes, providing an essential healthy baseline to study network rewiring in development and disease..

By mapping the GR network across multiple lineages, our data exposes endogenous features of the GR complex that offer physical rationale for long-standing clinical paradoxes in steroid therapy. A primary example is the tissue-restricted recruitment of the SMARCD3 (BAF60c), one of three mutually exclusive SWI/SNF components associated with cell-fate determination^41–43^. While glucocorticoids are universally deployed to suppress inflammation, their concurrent impact on solid tumour tissue remains double-edged, being pro-tumourigenic in TNBC yet protective in ER+ contexts. Our characterisation that GR physically engages SMARCD3 exclusively in epithelial lineages provides a molecular model for the pleiotropic effects of glucocorticoids. This result is clinically reflected in our survival stratification, where SMARCD3 expression mirrors the opposing trajectories of glucocorticoid signalling in breast cancer subtypes. Recent work has established the core SWI/SNF complex as a gatekeeper for GR-driven transcriptional networks and proposed it as a promising therapeutic target for TNBC ^44^. We therefore hypothesise that this lineage specific assembly offers a therapeutic opportunity to protect TNBC patients by targeting the epithelial-restricted SMARCD3-GR while leaving systemic lymphoid-driven anti-inflammatory benefits fully intact.

DIANNeR reasserts the need for analysis of normal human derived tissues. Our results reveal that the GR interactome of MCF10A, one of the most widely used models for normal breast epithelium, fundamentally diverges from normal tissue. Mechanistically, this clustering is driven by a shift in coregulator preference, where native breast epithelium selectively engages epithelial differentiation factors GRHL1 and GRHL2, the MCF10A interactome has rewired to recruit the EMT drivers ZEB1 and ZEB2, aligning it more closely with the mesenchymal TNBC cell line MDA-MB-231. Crucially, because these interactomes are not confidently predicted by transcript expression alone, our findings underscore that GR signalling is altered in MCF10A cells due to a cryptic, partial basal-like phenotype confounded by the expression of epithelial markers^45^. These findings demonstrate the value that DIANNeR brings through bypassing cell-line models for the study of transcription factor regulatory networks within normal tissue.

We identified a FOXP3–BCL11B–GR network in our primary CD4^+^ T cell population that was absent in the Jurkat model. Importantly, the FOXP3–GR interaction has been validated as specific to Treg cells in murine models ^46^, thus demonstrating a detection threshold capable of resolving sub-populations from a normal human sample. The failure of DDA-RIME to detect this FOXP3–BCL11B network in our benchmarking data supports our conclusion that DIANNeR overcomes the sampling limits of DDA-RIME based methods. This capability is critical because regulatory sub-populations, such as Tregs, can play a key role in the efficacy of glucocorticoid therapies^47,48^.

Finally, applying DIANNeR to the comparative analysis of normal human tissues, we demonstrate our method provides the necessary resolution to map changes in the GR network upon malignant transformation. For instance, while GR interactions with mesenchymal or EMT-associated factors diverge across model systems, the HOXA5–GR interaction remains preserved when comparing native normal breast epithelial cells to the non-tumourigenic MCF10A model, before being selectively lost in both MCF7 and MDA-MB-231 cell lines. Forced genetic expression of HOXA5, or its upregulation via retinoid therapy has been shown to induce apoptosis in MCF7 cells^49^ and impedes tumour initiation and progression by enforcing epithelial traits^50,51^. However, a persistent caveat of these studies has been their reliance on MCF10As as a proxy for normal tissue architecture presenting, based on our own findings, an important opportunity for follow-up studies in non-immortalised models. Crucially, by mapping primary ER+ breast cancer patient-derived xenograft models we provide evidence to support the loss of the HOXA5–GR axis *in vivo*. Mirroring this specificity, the absence of the Treg-associated BCL11B–FOXP3–GR axis in our PDX cohorts reinforces that DIANNeR discriminates genuine tissue-restricted pathology from overlapping physiological backgrounds

In conclusion, DIANNeR brings discovery-scale TF interactomics to scarce, non- transformed primary cell types for the first time. By establishing RIME for these sample types, we were able to resolve previously unmapped context-specific networks across health and disease, establishing a critical baseline with the potential to predict and mitigate adverse pleiotropic effects.

## Online Methods

### Cell culture

We analysed normal cells from three tissues, breast, blood and ureter, alongside five matched cell lines. For each cell line, three biological replicates were generated from independent passages. In the case of normal cells, each biological replicate was obtained from a different donor. For our analysis of normal human breast epithelial^52^ and NHU^53,54^ cells, we performed three biological replicates; and for our normal CD4^+^ T lymphocyte samples, we undertook four biological replicates.

All cell types were cultured at 37 °C in a humidified atmosphere containing 5% carbon dioxide in air. MCF7, MCF10A, MDA-MB-231, KMBC2 and Jurkat cell lines were routinely tested and confirmed negative for Mycoplasma spp. infection using Mycostrip (rep-mys-20, Invivogen). The authenticity of these five established human cell lines was confirmed by short tandem repeat (STR) profiling. Primary human breast epithelial cells were obtained from the Breast Cancer Now Biobank (REC 21/EE/0072). Breast tissue from cosmetic reduction mammoplasty was digested using a two-step enzymatic process, collagenase and hyaluronidase treatment, followed by sedimentation and subsequent trypsinisation to obtain a single-cell suspension. Breast epithelial cells were isolated using Fluorescence Activated Cell Sorting (FACS) using anti-EPCAM PE (347198, BD biosciences). Sorted cells were then cultured on collagen-coated plates and expanded with up to 2 passages to reach the number of cells required. Full details are provided in references ^52,55,56^ and Supplementary Materials. Cells were frozen at 5–10 × 10^6^ cells per vial and were shipped on dry ice to the University of York. Human ureter specimens were collected for research under NHS Research Ethics Committee approval Leeds (East) REC 99/095 for anonymous use of surplus tissue following renal transplant surgery. Normal human urothelial (NHU) cells were isolated for primary cell culture from surgical tissues as described previously ^54^. For our study, NHU cell culture was initiated by thawing a vial of cryopreserved cells into one T-75 flask. The cells were maintained in KSFM (Thermo Fisher) complete medium. Cultures at passage 4 were used for RIME assays. Human blood samples were obtained from the York Tissue Bank Biofluid service, University of York. Primary CD4^+^ T lymphocytes were isolated by immunomagnetic negative selection using an EasySep Human CD4^+^ T Cell Isolation Kit (17952, STEMCELL Technologies) and an EasySep Magnet (18000, STEMCELL Technologies). CD4^+^ T cell cultures were initially seeded at 1 × 10^6^ cells/mL in the culture medium. The time in culture was restricted to a maximum of 7 days to prevent T cell differentiation. Purity was assessed by flow cytometry panel **(Table S2)**. Subject to achieving the desired cell count, all non-immortalised normal cells in finite culture, as well as established cell lines, were treated with 100 nM dexamethasone for 1 hour before harvest. Detailed cell culture methods are provided in the following subsections.

### Breast tissue preparation

Tissue obtained from patients undergoing cosmetic reduction mammoplasty surgery is washed once with 70% ethanol and then three times with RPMI-1640 medium (Sigma R5886) containing 25 mM HEPES, supplemented with 5% fetal bovine serum (FBS, Gibco), 100U/ml penicillin, 0.1 mg/ml streptomycin (pen/strep Sigma P4333), 5 µg/ml amphotericin-B (Fungizone Sigma A2942) and 20 μg/ml Gentamicin (Sigma G1397).

Tissue is chopped into small pieces and digested for 12 to 16 hours at 37°C on a rotary shaker in RPMI-1640 medium plus 25mM HEPES, supplemented with 5% FBS, pen/strep and 5 µg/ml amphotericin-B containing 1mg/ml collagenase 1A (Sigma C2674) and hyaluronidase 1-S (Sigma 1-13506). The digested tissue is centrifuged at 380g for 20 minutes and washed in medium three times to remove enzymes. The tissue isolates are then sedimented three times at 1 × *g* for 30 minutes to collect the denser ductal tree fragments (TDLUs and ducts). The ductal tree fragments (DTF) are centrifuged (380 × *g*, 3 minutes) and frozen in medium containing 40% FBS and 6% DMSO (Sigma D2650) and stored in liquid nitrogen for later specific cell isolation.

### Isolation and expansion of luminal epithelial cells

The ductal tree fragments are digested using 0.25% trypsin/0.1% EDTA (Fisher Scientific Hyclone SV30031.01) and DNAse I (Boehringer/Roche Diagnostics now Sigma (Merck) 10104159001) is added to ensure the cells are single. Cells are counted before adding fluorescently conjugated antibodies to EPCAM (Becton Dickinson FITC 347197) and CD10 (Becton Dickinson APC 332777) and then sorted using Fluorescence Activated Cell Sorting (FACS) to separate the EPCAM+ cells (epithelial cells), CD10^+^ cells (myoepithelial cells) and double negatives (microenvironment cells). The fraction containing epithelial cells is then centrifuged (380 × *g*, 3 minutes) and the cell pellet resuspended, counted and cultured in DMEM:F12 containing 15 mM HEPES (Sigma D8437), supplemented with 10% FBS, 0.5 µg/ml hydrocortisone (Sigma H0888), 10 µg/ml apo-transferrin (Sigma T1147), 5 µg/ml insulin (Sigma I9278), 10 ng/ml EGF (Sigma E9644), pen/strep and 2.5 µg/ml amphotericin-B (Breast Culture Medium, BCM) on collagen coated 6 well tissue culture plates. Epithelial cells are grown for 1 passage before being frozen down in epithelial cell cryofreezing medium (BCM containing 40% FBS and 6% DMSO) and stored in liquid nitrogen.

For this work, vials of 300,000 passage 1 epithelial cells were thawed and seeded into collagen-coated 6-well tissue culture plates at a density of 100,000 cells per well. Epithelial cells were grown in BCM to passage 2 then trypsinised off the 6-well plates and seeded into collagen-coated T75 tissue culture flasks at a density of 1 x 10^6^ per flask for further expansion in BCM to passage 3. Once the cells had reached confluency, the medium was changed from BCM to a medium without growth factors, DMEM:F12 containing 15 mM HEPES, supplemented with 10% FBS, pen/strep and 2.5 µg/ml amphotericin-B (Fibroblast Medium, FM). After 1 day in this medium, the cells were treated with FM containing 100 nM dexamethasone for 1 hour. At this point, trypsinised cells were counted and either transferred to Eppendorf tubes, centrifuged at 12,000 rpm for 2 minutes using a microfuge or resuspended in epithelial cell cryofreezing medium and transferred to cryovials at a density of 5-10 x 10^6^ cells/vial. Dry cell pellets and cell suspensions in cryofreezing medium were then frozen at −80°C until further use. Using this technique, epithelial cell numbers of up to 140 x 10^6^ cells were achieved.

### Culture of normal human urothelial (NHU) cells

Normal human urothelial (NHU) cells were isolated for primary cell culture from surgical tissues and serially maintained as finite lifespan cell lines as described (Southgate *et al.*, Lab Invest. 1994). For our study, NHU cell culture was initiated by thawing a vial of cryopreserved cells into one T-75 flask. The cells were grown to near confluence in KSFM (Thermo Fisher) medium containing recombinant human epidermal growth factor (5ng/mL, Gibco), bovine pituitary extract (50 ng/mL, Gibco), and cholera toxin (30ng/mL), herein referred to as KSFM complete medium (KSFMc). Upon reaching near confluence, the cells were split 1:3 and subcultured in T-75 flasks, continuing their growth in the KSFMc medium. When confluence was achieved, cells were divided into four 15 cm dishes using approximately a 1:3 split ratio, and the cultures were maintained in KSFMc. Cultures at passage 4 were treated with 100nM dexamethasone and crosslinked for RIME assays.

### Blood collection and isolation of peripheral blood mononuclear cells (PBMCs)

Human blood samples were obtained from the York Tissue Bank biofluid service, University of York. After obtaining written, informed consent (patient information sheet and consent form, version 3.5 or earlier, approved by the Biology Ethics Committee, University of York, reference number IH202201), venous blood was collected from healthy adult human volunteers into vacutainers containing the active anticoagulant trisodium citrate with citric acid and dextrose, solution A (366645, Fisher Scientific). Samples were provided pseudonymised to the researcher and processed within 1 hour upon blood collection.

Whole blood was transferred from the collection tube to a sterile 50 mL Falcon tube (62.547.254, Sarstedt) using a disposable, sterile Pasteur pipette. The blood was then diluted at a 1:1 ratio with sterile PBS. To prepare the Leucosep tube for gradient separation, 15 mL of Lymphoprep reagent (07801, STEMCELL Technologies) was added, and the tube was briefly centrifuged to ensure proper positioning of the reagent. The diluted blood was carefully layered onto the Lymphoprep in the Leucosep tube, which was then centrifuged at 800 x *g* for 15 minutes at 25°C. The buffy coat containing PBMCs was gently aspirated using a sterile Pasteur pipette and transferred to a new sterile 50 mL Falcon tube. The volume was brought up to 50 mL with sterile PBS, and the tube was centrifuged at 250 x *g* for 10 minutes at room temperature to pellet the PBMCs. Following centrifugation, the supernatant was discarded, and the PBMC pellet was resuspended for immediate isolation of CD4^+^ T cells.

### CD4^+^ T lymphocyte isolation and *in vitro* expansion

Human blood samples were obtained from the York Tissue Bank biofluid service, University of York. Upon receipt, blood samples were immediately processed. Primary CD4^+^ T lymphocytes were isolated by immunomagnetic negative selection using an EasySep Human CD4^+^ T cell Isolation Kit (17952, STEMCELL Technologies) and an EasySep Magnet (18000, STEMCELL Technologies). PBMCs were resuspended at 5 x 10^7^ cells/mL in EasySep buffer (2% FBS and 1 mM EDTA in sterile PBS, Ca^2+^ and Mg^2+^ free). The isolation process followed the manufacturer’s protocols. CD4^+^ T cell cultures were initiated at 1 x 10^6^ cells/mL in the culture medium (DMEM, 1X L-Glutamine, 1 mM Sodium Pyruvate, 10% FBS, 50 µM beta-mercaptoethanol, 1X MEM non-essential amino acid solution, 1X pen/strep). To promote *in vitro* proliferation, T cells were activated using 25 μL/mL of ImmunoCult Human anti-CD3/CD28/CD2 T Cell Activator (10970, STEMCELL Technologies) and 20 ng/mL recombinant human IL-2 (rh-IL-2, 791904, BioLegend). Three days after the initial seeding, cultures were expanded to a final density of 1 x 10^5^ to 2.5 x 10^5^ cells/mL, with rh-IL-2 (20 ng/mL) added accordingly. This expansion was repeated on day 5 and, if necessary, on day 7. Subject to achieving the desired cell count, cells were treated with 100 nM dexamethasone before fixation with formaldehyde. As prolonged culture may lead to T cell differentiation, we restricted the *in vitro* culture to a maximum of 7 days.

### Flow cytometry

To assess the purity of isolated CD4^+^ T cells, immunophenotyping was performed by flow cytometry. For each staining condition, 10,000 PBMCs or isolated CD4^+^ T cells were transferred into a conical 96-well plate (83.3926500, Sarstedt). Cells were washed once with PBS and resuspended in 100 μL of FACS buffer (2% FBS in sterile PBS). The appropriate antibody panel (**Supplementary Table S2**) was added directly to the cell suspension, followed by incubation at 4°C for 20-30 minutes in the dark. Cells were washed twice with 100 μL of FACS buffer or PBS and resuspended in 100 μL of FACS buffer for analysis.

Flow cytometry was performed using CytoFLEX or CytoFLEX S cytometers (Beckman Coulter). Data was analysed with CytExpert software. Live, single cells were gated based on forward and side scatter (FSC/SSC) properties. Leukocytes were identified as CD45^+^, and monocytes excluded as CD14^+^ T cells were gated as CD3^+^ within the CD45^+^ CD14^-^ population. CD4^+^ T cells were identified as CD3^+^CD4^+^CD8^-^. The proportion of CD4^+^ T cells was quantified to assess isolation purity. Gating boundaries were established using PBMCs stained with all relevant antibodies as positive controls for each CD marker, and live/dead discrimination was calibrated using cells heat-killed at 65 °C for 15 minutes.

### Patient Derived Xenografts (PDX)

All *in vivo* studies were carried out in accordance with the UK Home Office (Scientific Procedures) Act 1986 under project licence PPL40/3645 and as described by Simões *et al.* ^57^. Analytical material was derived from three distinct xenografts (BB3RC31, BB3RC29 and HBCx34 ^57,58^). HBCx34 PDX was established from a luminal B breast cancer sample as previously described by Cottu *et al.* ^58^. In summary, small fragments of PDX tumours were implanted subcutaneously into the flanks of 8–12-week-old female NSG (NOD scid gamma, NOD.Cg Prkdcscid Il2rgtm1Wjl/SzJ) mice. These preclinical models are estrogen-dependent, so animals were administered 8 μg/ml 17-beta estradiol (Sigma-Aldrich, #E2758) in drinking water at least 4 days prior to implantation until the end of the experiment.

### Culture of established cell lines

MCF7 cells were cultured in the DMEM (21969-035, Gibco) supplemented with 10% FBS (Gibco) and 1% L-glutamine. Cultures were maintained in a T-75 flask (Corning) before being seeded into a 15 cm Petri dish (Corning). Cultures were harvested for the RIME protocol at 90% confluence, providing approximately 20 million cells per dish. MDA-MB-231 cells followed the same culturing protocol as MCF7 cells. KMBC2 followed the same protocol with an altered cell culture medium consisting of 50% DMEM (21969-035, Gibco), 50% RPMI (31870-025, Gibco) and 5% FBS (Gibco). Jurkat cells were cultured per ATCC’s handling procedures, being initiated at a low cell density of 1-2 x 10^5^ cell/mL in RPMI medium supplemented with 10% FBS. Following the initial lag phase, cell density was maintained between 1 x 10^5^ to 1 x 10^6^ viable cells/mL. Subculturing of Jurkat cells included pre-warming the fresh growth medium in a CO_2_-controlled cell culture incubator for 15 to 20 minutes to equilibrate temperature and pH, ensuring optimal cell viability.

### Optimisation of GR Antibody

To capture the acute GR nuclear interactome, cells were treated with 100 nM Dexamethasone (Dex) for 1 h prior to harvesting. ^59,60^. Three candidate antibodies (Atlas HPA004248, CST 12041, and Thermo PA1-511A) were benchmarked. The HPA004248 polyclonal antibody was selected based on superior bait coverage 38%, **Figure 1B**) and significant recovery of known interactors from BioGRID (*p* = 6.07 × 10^-22^) and STRING (4.75 × 10^-34^) of these respective reference datasets. Specificity was further confirmed via enrichment analysis (gProfiler^61^); detected interactors showed significant overrepresentation of nuclear and chromatin-associated GO terms, with corresponding depletion of cytosolic proteins (*p =* 1.09 × 10^−5^).

### Rapid Immunoprecipitation Mass spectrometry of Endogenous proteins (RIME)

Our DIA-NN-enabled RIME (DIANNeR) protocol builds on the established RIME methods^1,2,62^, with specific modifications tailored to enhance performance across normal cells in finite culture. For each condition, a minimum of three biological replicates were required. For normal cells in finite culture, biological replicates were obtained from three independent donors; for established cell lines, biological replicates were defined as separate passages. For PDXs, we analysed one technical replicate from independent PDX tumours to provide donor-level biological replication consistent with our approach for normal cells. The following steps are provided as an experimental protocol on protocols.io, [Reviewer link].

Adherent cell cultures (MCF7, KMBC2, MDA-MB-231, normal breast epithelial and NHU cells) were subjected to the protocol as follows, which is optimised for 15 cm plates. If higher cell counts are required, multiple plates can be processed in parallel and the lysates combined after sonication. For each plate, the medium was removed and cell culture washed with 20 ml ice cold PBS before fixation. For each 15 cm dish, adherent cells were treated with a 20 ml solution of 2 mM disuccinimidyl glutarate (DSG, sc-285455A, Santa Cruz Biotechnology) in PBS for 20 minutes at ambient temperature. Subsequently, the DSG solution was discarded, and the cells were further crosslinked for 10 minutes using a 1% formaldehyde solution (28908, Thermo) in PBS at ambient temperature. Quenching was achieved by incubating the cells with 0.125 M glycine for 5 minutes. Following these steps, the crosslinking solution was removed and cells were washed twice with cold PBS and then harvested using a cell scraper and ice-cold PBS supplemented with 1X cOmplete Protease Inhibitor Cocktail (11873580001, Sigma). Cells were collected into 1.5 mL Protein LoBind tubes (EP0030108094, Eppendorf) and centrifuged (6000 × *g*, 4°C) for 3 minutes to pellet the cells and the supernatant was removed. Cells were then frozen at −70°C or processed immediately as per the sonication step.

For suspension (Jurkat and normal CD4^+^ T cells) culture, 16% formaldehyde was added directly to cultures to a final concentration of 1%. After crosslinking the cells for 10 minutes at ambient temperature, quenching was performed using 0.125M glycine for 5 minutes. Cells were pelleted by centrifugation (500 × *g,* 4°C) for 3 minutes, resuspended in ice-cold PBS supplemented with 1X cOmplete Protease Inhibitor Cocktail, and transferred to 1.5 mL Protein LoBind tubes. The cells were washed twice with PBS including protease inhibitor, supernatant removed, and either frozen at −70°C or processed immediately for sonication.

For sonication of non-PDX samples, PBS-washed cell pellets containing up to 20 million crosslinked cells were resuspended and incubated at 4°C in 1 mL of Lysis Buffer 1^1,2,62^ (composed of 50 mM HEPES, pH 7.5, 140 mM NaCl, 1mM EDTA, 10% glycerol, 0.5% NP-40, 0.25% Triton X100 in molecular biology grade water) for 10 minutes, followed by incubation in 1 mL of Lysis Buffer 2 ^1,2,62^ (10 mM Tris-HCl, pH 8.0, 200 mM NaCl, 1 mM EDTA, 0.5 mM EGTA in molecular biology grade water) for 5 minutes in sequence. After each incubation step, samples were centrifuged (2000 × *g,* 4°C) for 5 minutes to collect the cell/nuclei pellets. These pellets were then resuspended in 300 μL of Lysis Buffer 3, transferred to a 1.5 ml Bioruptor Pico Microtubes (Diagenode) and subjected to sonication with cycles of 30 seconds on / 30 seconds off using a Bioruptor Pico (Diagenode) for 5–10 cycles depending on the cell type and instrument. After sonication, 30 μL of the lysate was diluted with an equal volume of 1 × Tris/Glycine/SDS buffer (1610732, Bio-Rad) in 1:1 ratio. To confirm effective shearing, 10 µL of lysate was subject to RNAse A (2 μL) (AM2271, Thermo) treatment at 37°C for 1 hour, followed by proteinase K (2 μL) (10005393, Invitrogen) digestion at 65°C for 1 hour. The enzymatic reactions were terminated by heating at 95°C for 5 minutes. The purified DNA fragments were then visualised using SYBR Safe DNA Gel Stain (S33102, Thermo) on 1% agarose (A20090, Melford) gel with 1X TEA buffer as the running medium for gel electrophoresis. DNA fragments were visualised on a 1% agarose gel (Melford, A20090) stained with SYBR Safe DNA Gel Stain (Thermo, S33102) using 1X TAE buffer as the running medium. Samples that demonstrated adequate shearing (300–500 bp) proceeded to the next steps; otherwise, sonication conditions were optimised by adjusting the number of cycles. Following confirmation of adequate shearing, cell lysates were supplemented with Triton X-100 (X100, Sigma) to a final concentration of 1%. Subsequently, lysates were centrifuged (16,800 × *g*, 4°C) for 10 minutes to eliminate debris. The remaining supernatant was combined with previously prepared antibody-conjugated Pierce Protein A/G Magnetic Beads (88802, Thermo) and incubated overnight at 4°C with constant agitation. Depending on cell yield and experimental requirements, lysate from multiple 15 cm plates was pooled prior to the addition of the antibody conjugated beads.

PDX samples were provided frozen, these were fixed and cryosectioned without thawing to preserve integrity and enable the permeability of crosslinking reagents, adapting the method established by Papachristou *et al.* ^2^. PDX tissue samples were cryosectioned at 30 µm using a Leica CM1950 Cryostat. Sections were crosslinked in suspension with 2 mM DSG for 25 minutes, followed by 1% formaldehyde for 20 minutes. Crosslinking was quenched with 0.25 M glycine, and samples were centrifuged (2500 × *g*) for 3 minutes. Pellets were washed twice with cold PBS and resuspended in 6 ml LB3 ^1,2,62^ (composed of 10 mM Tris-HCl, pH 8.0, 100 mM NaCl, 1 mM EDTA, 0.5 mM EGTA, 0.1% sodium deoxycholate, 0.5% N-lauroylsarcosine sodium in molecular biology grade water) buffer, then subjected to sonication for 12–20 cycles, 30 seconds on / 1 minute off using a Ultrasonic Processor CP 750 with CV33 Probe (Cole Parmer) at 4°C, cycle time was based on tumour size and processed alongside other samples post-sonication.

For each RIME assay, 5 μg of anti-NR3C1 antibody (HPA004248, Atlas) or control normal rabbit IgG (2729, Cell signalling Technology) was employed. The bead-antibody conjugates were prepared by washing either 10 µL (< 10^7^ cells) or 50 µL (> 10^7^ cells or PDXs) of magnetic bead suspension three times with 1 mL of protein-free blocking buffer (37584, Thermo). The beads were then resuspended in 500 μL of the same buffer, and the appropriate antibody was introduced to the suspension. The mixture was incubated at ambient temperature for 1 hour or at 4°C overnight. To eliminate any non-specifically bound antibodies, the bead-antibody conjugates were washed 3 times with 1 mL of protein-free blocking buffer (37584, Thermo) before being combined with the cell lysates.

After the cell lysates were incubated overnight with bead-antibody conjugate at 4°C, the beads were subject to ten washes within a 4°C cold room with a washing buffer (composed of 50 mM HEPES pH 7.6, 1mM EDTA, 1% NP40, 0.5M lithium chloride, 0.7% sodium deoxycholate in molecular biology grade water). The beads were quickly rinsed twice with freshly prepared 100mM ammonium bicarbonate buffer, and stored and maintained at −70°C until mass spectrometry-based proteomics analysis. On the day of analysis, the RIME samples underwent on-bead trypsin digestion to produce peptides, following an established protocol ^1^.

### Mass spectrometry analysis

To minimise the effect of instrumental drift and technical variation, our DIANNeR samples were randomised and analysed concurrently using liquid chromatography (LC) followed by PASEF-DIA MS^63,64^. For our acquisition mode comparison, samples were run using a matched acquisition strategy. Samples were loaded onto a 8 cm Performance C_18_ column (8 cm x 150 µm, 1.5 µm), followed by 14.4 minute acquisition (100 SPD EvoSep One gradient) using a Bruker timsTOF HT in both DIA and DDA mode to provide a single experimental variable. All DDA and DIA comparisons utilised identical physical sample splits from each donor after RIME processing to eliminate biological and technical confounding. Additionally, samples were analysed using established LC and DDA acquisition methods for the Orbitrap Fusion Tribrid mass spectrometer to provide a comparative baseline. Full details of methods for our acquisition mode comparison are provided in Supplementary Methods.

For our multi-tissue analysis, DIA Sample acquisition order was randomised, using the RAND function in Microsoft Excel, to mitigate any potential bias resulting from time- or injection count-correlated performance changes. Following on-bead digestion, peptides were loaded onto EvoTip Pure tips for nanoUPLC using an EvoSep One system, followed by a pre-set 30 SPD gradient on a 15 cm EvoSep C_18_ Performance column (15 cm × 150 µm, 1.5 µm).

The nanoUPLC system was interfaced to a timsTOF HT mass spectrometer (Bruker) with a CaptiveSpray ionisation source. Positive PASEF-DIA, nanoESI-MS and MS^2^ spectra were acquired using Compass HyStar software (Bruker, version 6.2). Instrument source settings were: capillary voltage, 1,500 V; dry gas, 3 l/min; dry temperature; 180 °C. Spectra were acquired between *m/z* 100-1,700. DIA windows were set to 25 Th width between *m/z* 400-1201 and a TIMS range of 1/K_0_ 0.6-1.60 V.s/cm^2^. Collision energy was interpolated between 20 eV at 0.5 V.s/cm^2^ to 59 eV at 1.6 V.s/cm^2^.

The resulting LC-MS data (acquired in either Thermo’s proprietary RAW format or Bruker’s proprietary .d format) was processed using either FragPipe (v23)^65^ for DDA data or via DIA-NN (2.0) software, searching against a human-specific SwissProt database appended with sequences of common proteomic contaminants. For the DIA pipeline, an *in-silico* predicted spectral library was created with the DIA-NN software, and subsequently refined through iterative searches against the DIA datasets resulting from the samples. Search criteria for both workflows were set to maintain a false discovery rate (FDR) of 1%. Protein-level quantification was extracted using the high-precision quant-UMS tool within DIA-NN^66^, culminating in the production of a .parquet output. This output was subsequently processed using custom KNIME workflows to yield a protein group-centric dataset with additional filtering to protein (q < 0.01, minimum of 2 peptides). Finally, downstream differential enrichment analysis for both the DIA and DDA workflows was undertaken using the FragPipe-Analyst pipeline^5^ with Limma to establish log_2_ fold changes (LFC) and p-value for specific interactors, with sample minimal imputation and p-values subjected to Hochberg-Benjamini correction.

The NR3C/GR sequence coverage plots for the comparison of antibodies were generated by the Scaffold Proteome Software.

### DIA Database Searching

LC-MS data, in Bruker .d format, was processed using DIA-NN (v2.0) software and searched against an *in-silico* predicted spectral library, derived from the human subset of the SwissProt database appended with common proteomic contaminants. Search criteria were set to maintain a false discovery rate (FDR) of 1% and specified the following parameters: --qvalue 0.01 --matrices --gen-spec-lib --met-excision --cut K*, R* --missed-cleavages 1 --individual-mass-acc --individual-windows --no-prot-inf --reanalyse --rt-profiling --no-norm. Peptide-centric output, in .tsv format, was pivoted to protein-centric summaries using KNIME (v5.1) and data filtered to require protein q-values < 0.01 and a minimum of two peptides per accepted protein.

### Quantitative DIA-NN MS multi-tissue dataset analysis

The GR DIA-NN-enabled RIME .tsv output was processed using the publicly hosted FragPipe-Analyst pipeline (http://fragpipe-analyst.nesvilab.org/) into .RData form and then analysed using the FragPipeAnalystR package.

### GO, Reactome, and KEGG over-representation analysis

Specific interactions were derived from differential enrichment analysis comparing the GR IP group to the IgG control group (LFC > 2, *p*-adjust < 0.05). Overrepresentation analysis of multiple terms was performed against GO, Reactome and KEGG terms through the g:Profiler (https://biit.cs.ut.ee/gprofiler/gost) R package^61,67^ with a p-adjust threshold of 0.05 Overrepresentation of specific terms was performed using clusterProfiler (v4.12.6)^68^ and the *enricher* function.

### Transcriptomic data correlation analysis

Expression data for each GR-interacting protein was obtained for luminal cells from normal mammary epithelium, urothelial cells from the normal ureter, and CD4^+^ T cells from normal blood from across more than 40 publicly available datasets accessed via the CellxGene API^69–75^. Integrating > 40 lineage-matched single-cell references provided a robust, conservative test of global predictability/ As DIANNeR reports bulk interaction abundances, single-cell counts were pseudobulked on a per-dataset basis and averaged to generate mean counts-per-ten-thousand (CPTT) values for each cell population. cDNA libraries for cancer cell lines were generated from matched stocks of cells to those used for the DIANNeR dataset, and sequenced in isogenic triplicate, and transcript per million (TPM) values were averaged across replicates. To harmonise across sequencing platforms and quantification methods, expression measures were transformed to z-scores before Pearson’s correlation analysis **(Table S3)**.

### ChIP-seq colocalisation analysis

Colocalisation between GR and other transcription factors was assessed using ChIP-Atlas, which reports peak colocalisation across uniformly processed public ChIP-seq datasets^76^ aligned to H. sapiens genome assembly hg38. We queried NR3C1 (GR) ChIP-seq experiments against ChIP-seq datasets for other transcription factors. The resulting overlap matrices were downloaded as CSV files, imported into R, and filtered to retain only samples related to the cell types and transcription factors of interest **(Table S4-8)**. No additional peak calling or reprocessing was performed; all colocalisation values reflect the standardised ChIP-Atlas pipeline.

### Survival Data

Patient survival and clinical correlation analyses were performed using the KM Plotter breast cancer database (kmplot.com). The cohort was restricted by receptor status on the basis of immunohistochemistry (IHC) for ER and microarray for HER2 criteria. Redundant samples and biased arrays were excluded using the integrated pipeline. To determine the prognostic threshold, patients were stratified into high and low expression cohorts using the integrated percentile cutoff algorithm, calculated over the entire database.

## Supplementary Data

Supplementary data are available online.

## Data Availability

All proteomic mass spectrometry datasets and results files are deposited in ProteomeXchange (PXD066547) and available to download from MassIVE (MSV000098634) [doi:10.25345/C5707X199]. Bulk RNA-seq data supporting Figure 8 have been deposited in ArrayExpress under accession number E-MTAB-16514. The processed data for GR interactome across tissues is provided in supplementary file **Results File S1,** and via Shiny app at https://holding-lab.shinyapps.io/DIANNeR-Volcano/, doi:10.5281/zenodo.21415678. The R code for data analysis and figures is available from https://github.com/Holding-Lab/DIANNeR, doi:10.5281/zenodo.16762025.

## Ethical Approval

This research was approved by the Biology Ethics Committee (BEC), University of York (AH202111).

## Supporting information

Results File S1

## Acknowledgements

The authors would like to thank all the anonymous volunteers from the University of York for their blood donation and the Biofluid service phlebotomists. The NHU cell culture was supported by a programme grant to the Jack Birch Unit from York Against Cancer. Jurkat cells were a kind gift from Dr Dimitris Lagos Lab, University of York. The authors wish to acknowledge the role of the Breast Cancer Now Biobank in collecting and making available the samples used in the generation of this publication, and all the patients who donated the samples. The authors would like to thank the Bioscience Technology Facility, University of York for their support in mass spectrometry analysis and flow cytometry analysis. The York Centre of Excellence in Mass Spectrometry was created thanks to a major capital investment through Science City York, supported by Yorkshire Forward with funds from the Northern Way Initiative, and subsequent support from EPSRC (EP/K039660/1; EP/M028127/1). The Viking cluster, which is a high-performance compute facility provided by the University of York, was used during this project. We are grateful for computational support from the University of York, IT Services and the Research IT team.

## Extended Data Figures

**Extended Data Figure 1.**
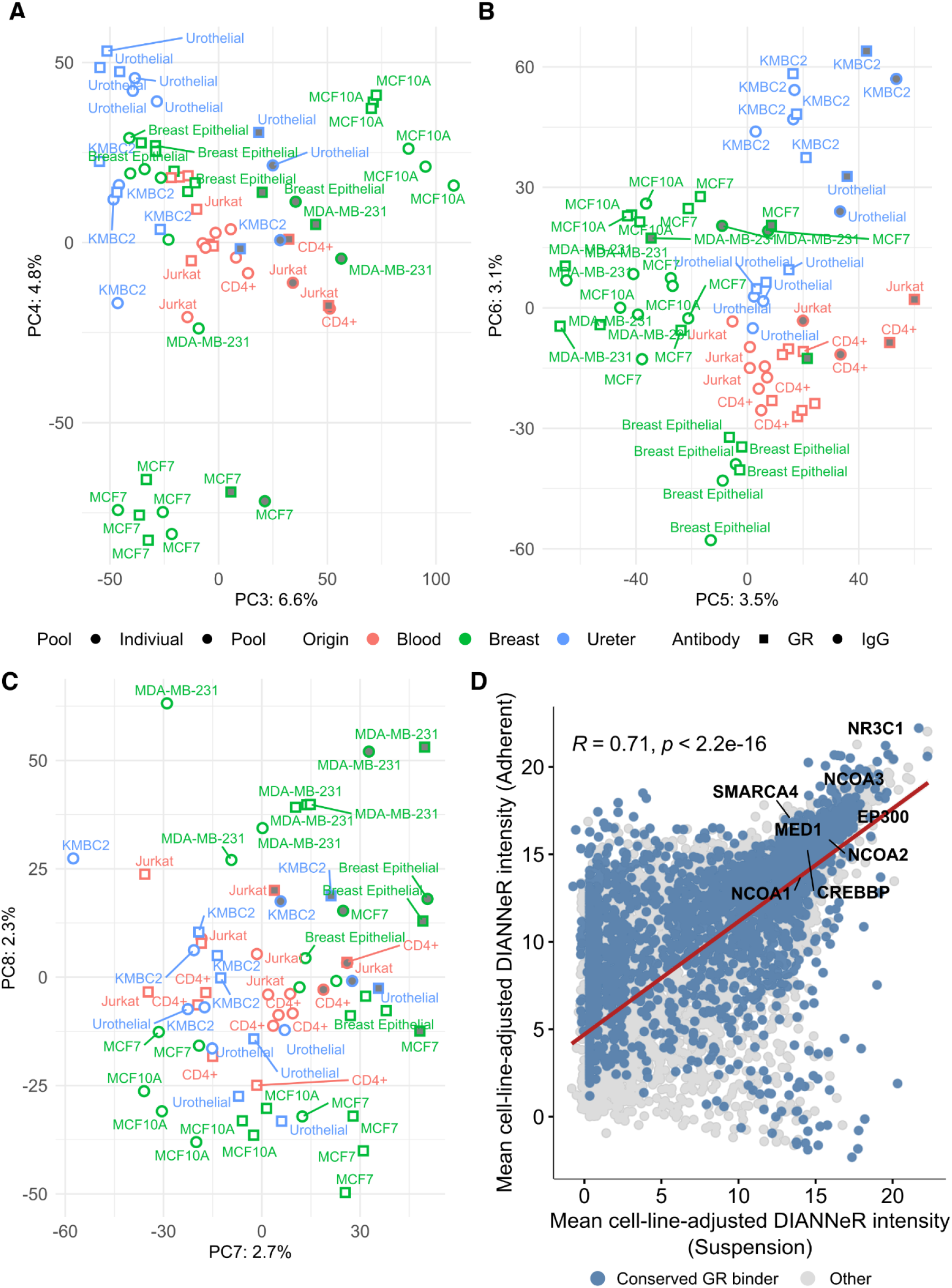
Visualisation of higher order principal components (PC3–PC8) and analysis of the residual effects associated with culture format and processing. **(A)** PC3 captures variance associated with the MCF10A samples, and PC4 isolates MCF7 samples. **(B)** PC5 separates the breast cancer cell lines from normal epithelial samples and from blood and ureter. PC6 is specific for differentiating the KMBC2 samples and Breast Epithelial samples from the other tissues analysed. **(C)** PC8 distinctly captures the MDA-MB-231 cells, separating them from MCF10A and MCF7. **(D)** Protein-level GR-associated intensities were adjusted for cell-line-specific effects using linear modelling while preserving adherent vs suspension culture and process effects. Mean adjusted intensities were compared between suspension- and adherent-derived systems. Each point represents a protein. Canonical GR cofactors are highlighted. A strong positive correlation (Pearson R = 0.71, p < 0.0001) indicates a high correlation of GR interactive measurements irrespective of culture format, supporting that the observed experimental differences are driven by the underlying biology.

**Extended Figure 2.**
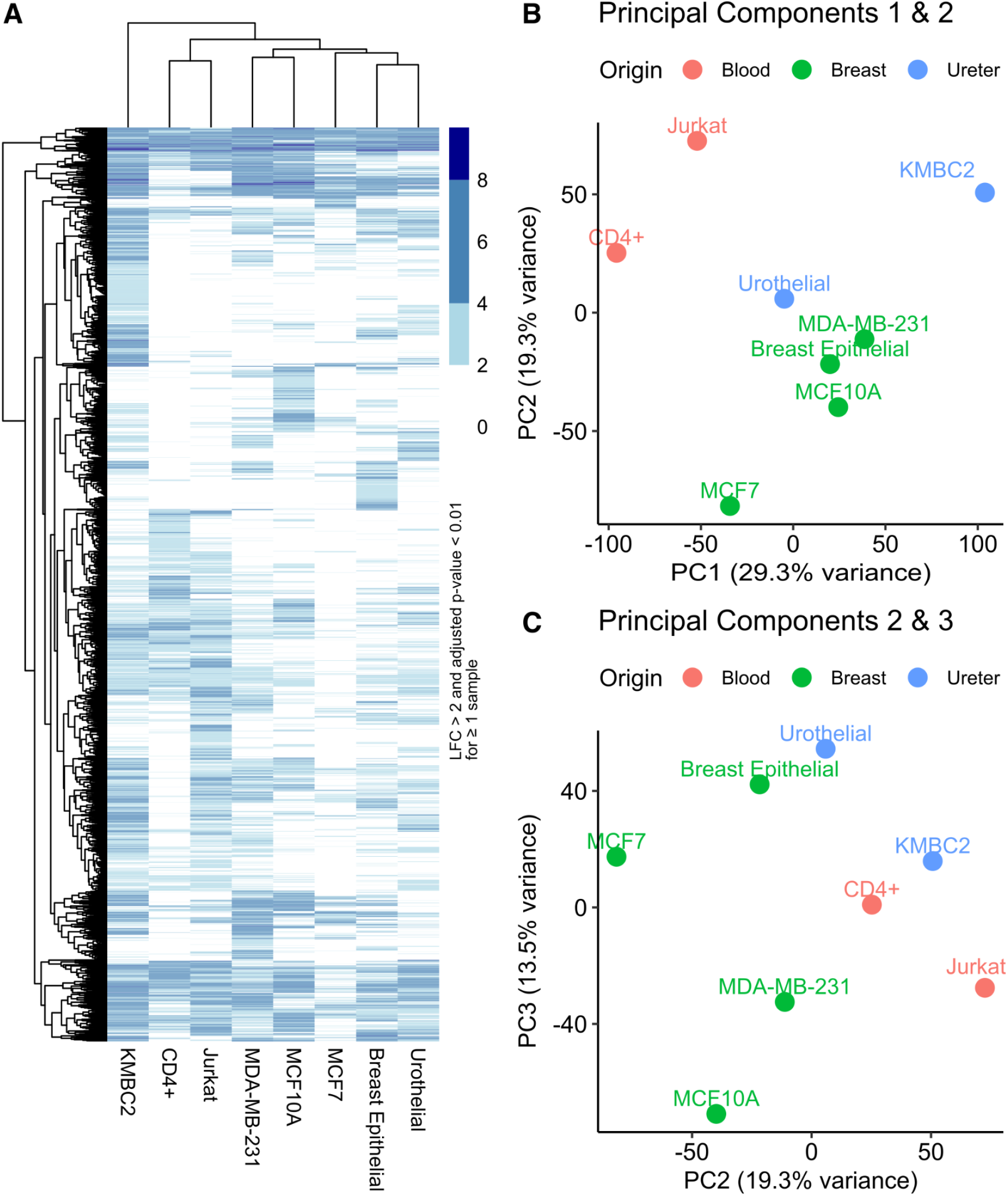
The GR interactome in normal CD4+ T-cells and Jurkat cells is distinct from that of epithelial cells, and GR interactions in the normal breast epithelium and urothelium are more similar than both tissues are to their matched cell lines. **(A)** Clustering of all detecting GR interacting proteins (LFC > 2 and p-value < 0.01 in ≥ 1 sample) by LFC shows similar clustering to total protein intensity. Jurkat and CD4+ T-cells clustered together, as do the breast epithelial and urothelial interactomes. KMBC2 was the only sample that displayed different clustering between the complete dataset and the prioritised heatmap, with the KMBC2 GR interactome clustering more closely to that of MDA-MB-231 and MCF10A, whereas in the full dataset, the KMBC2 data clustered separately. **(B)** PCA clustering by PC1 and PC2 of GR interactomes. **(C)** PCA clustering by PC2 and PC3 of GR interactomes.

**Extended Data Figure 3.**
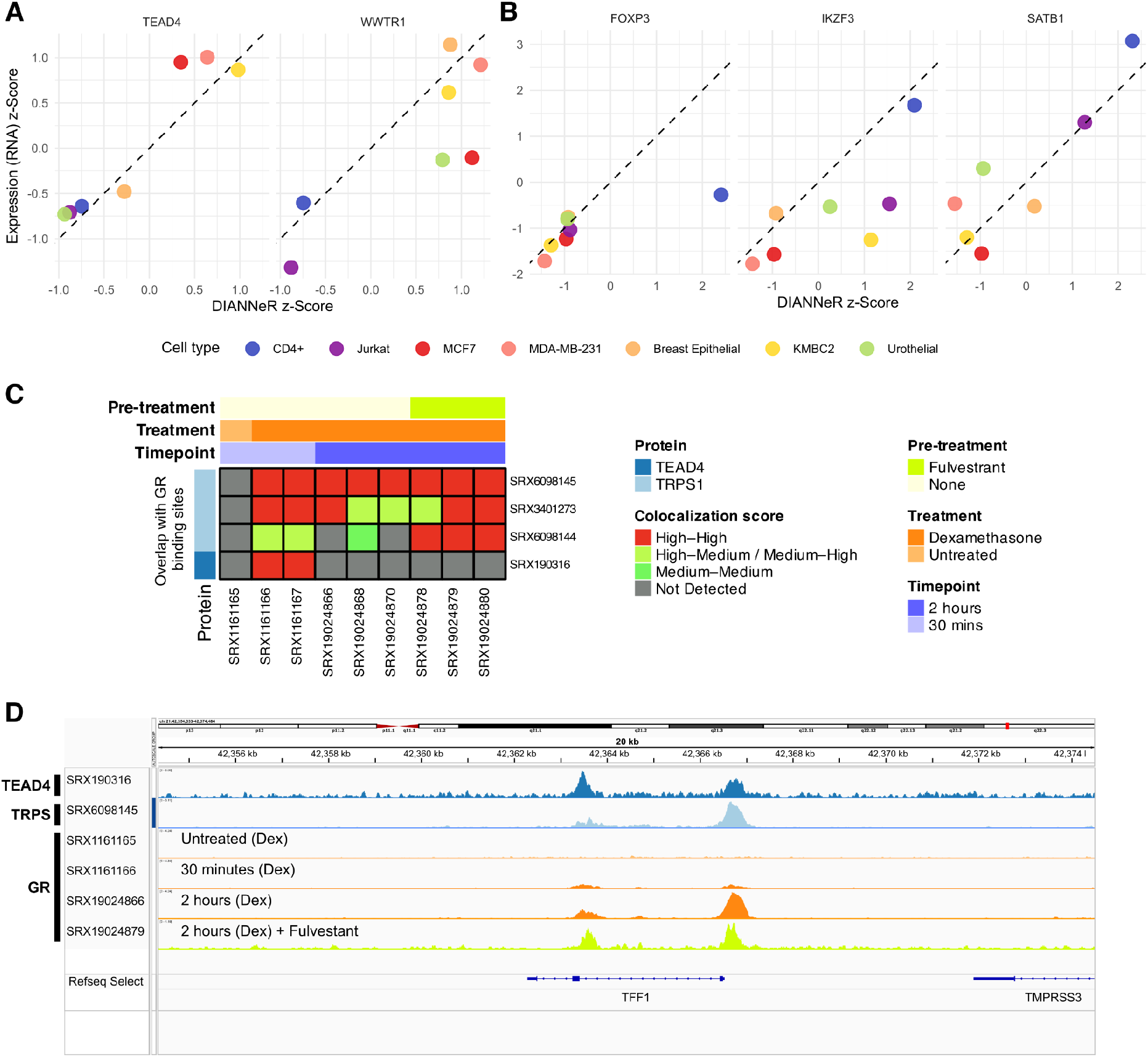
**(A)** Correlation plot between transcript expression (z-Score normalised) and DIANNeR LFC (z-Score normalised) for GR interacting proteins TEAD4 and WWTR1, also known as TAZ **(B)** Correlation plot between transcript expression (z-Score normalised) and DIANNeR LFC (z-Score normalised) for GR interacting proteins FOXP3, IKZF3 and SATB1. **(C)** ChIP-Atlas colocalisation analysis of TRPS–GR and TEAD4–GR binding sites in MCF7 cells. **(D)** IGV browser screenshot for ChIP-seq tracks for GR, TEAD4, and TRPS1 binding in MCF7 cells at the TFF1 locus.

