## Supplementary Materials for "DIANNeR: label-free RIME resolves lineage-restricted Glucocorticoid Receptor interactomes in normal human tissues"

Weiye Zhao**[1], Thomas F Grimes**[1], Susanna F Rose[1], Jack Stenning[1], Chris Taylor[2], Chloë Baldreki[2], Iain Goulding[4], Ros Duke[1], Simon C Baker[1], Marcela Montes de Oca[1,3], Aparna D. Sinha[1], Jenny Hinley[1], James M Fox[1], Paul M Kaye[3], Jennifer J Gomm[4], Louise J Jones[4], Elisabetta Marangoni[5], Bruno M Simões[6], Robert B Clarke[6], Jennifer Southgate[1], Katherine S Bridge[1], Adam Dowle[2], Andrew N Holding[1*]

1. Department of Biology and York Biomedical Research Institute, University of York, York YO10 5DD, UK
2. Metabolomics and Proteomics, Biosciences Technology Facility, Department of Biology, University of York, York YO10 5DD, UK
3. Hull York Medical School and York Biomedical Research Institute, University of York, York YO10 5DD, UK
4. Breast Cancer Now Biobank, Centre for Tumour Biology, Barts Cancer Institute, John Vane Science Centre, Queen Mary University of London, London EC1M 6AU, UK
5. Translational Research Department, Institut Curie, 26 rue d'Ulm, 75005 Paris, France
6. Manchester Breast Centre, Division of Cancer Sciences, Faculty of Biology, Medicine and Health, University of Manchester, Manchester, UK

** These authors contributed equally to this work.

### Supplementary  Figure 1

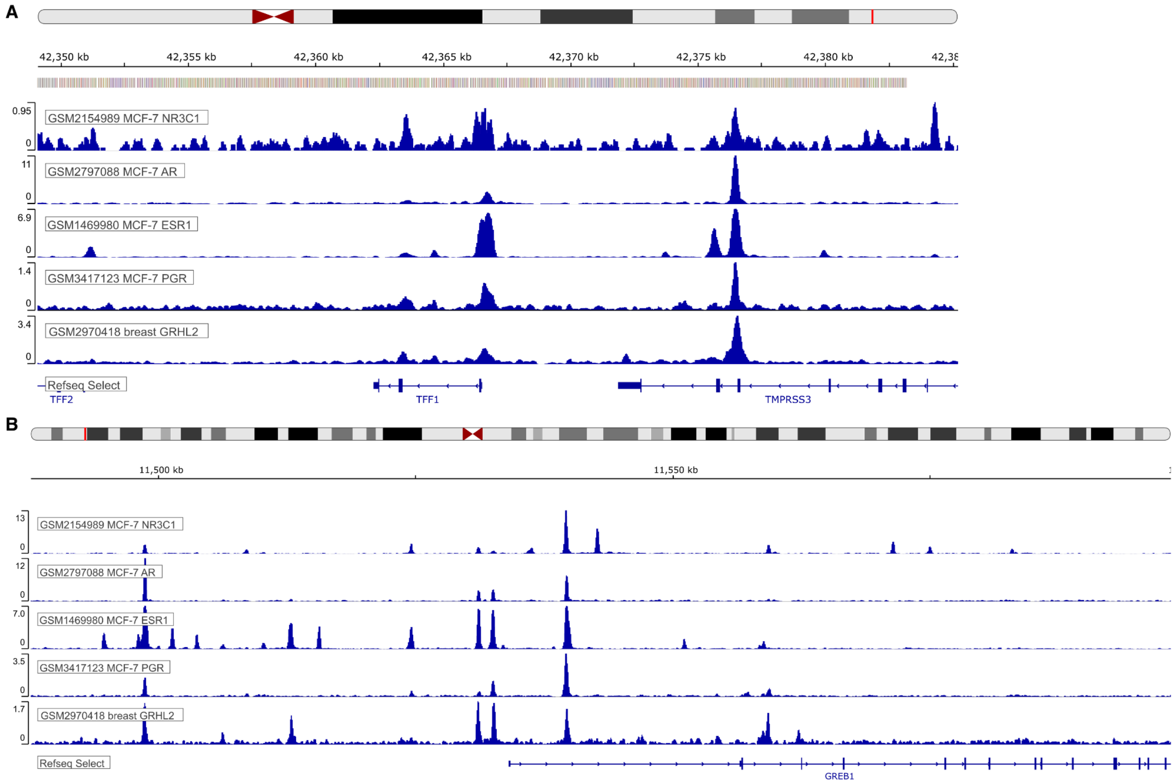


**Supplementary Figure 1** Exported ChIP-seq track view from Cistome.Org (Taing et al., Nucleic Acids Res., 2024) **(A)** for GR, AR, ERα, PR and GRHL2 binding at TFF1 and **(B)** for GR, AR, ERα, PR and binding GRHL2 at GREB1 loci. Individual tracks are labelled with GSM reference on the figure.

### Supplementary Figure 2

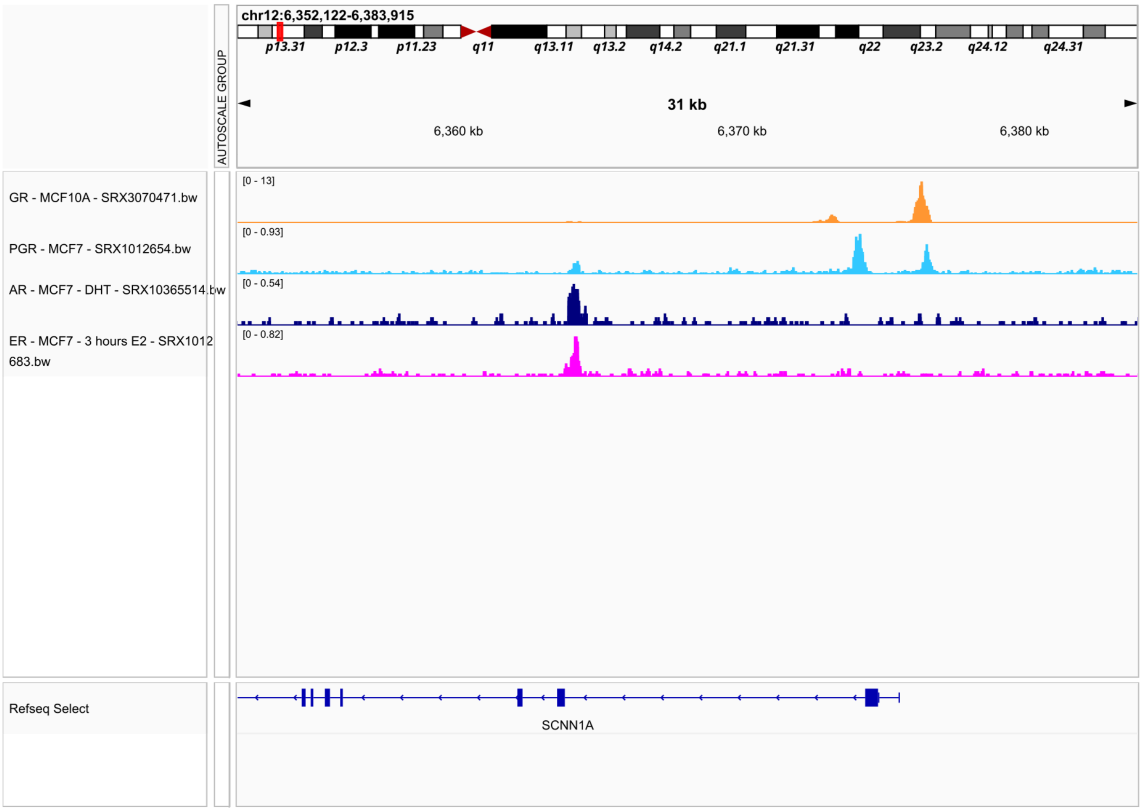


**Supplementary Figure 2** IGV browser screenshot for ChIP-seq tracks for GR binding in MCF10A and PR, AR and ERα in MCF7s at the SCNN1A locus. Individual tracks are labelled with SRX reference on the figure.

### Supplementary Tables

#### Table S1

| **Model** | **Tissue** | **Morphology** | **Status** | **Notes** |
|---|---|---|---|---|
| Normal Breast Cells | Breast | Epithelial | Normal | Finite Culture<br>Not immortalised |
| Normal Urothelial Cells (NHU) | Bladder | Epithelial | Normal | Finite Culture<br>Not immortalised |
| Normal $CD4^+$ T-cells | Blood | Lymphocyte | Normal | Finite Culture<br>Not immortalised |
| MCF7 | Breast | Epithelial | Carcinoma | ERα+ Breast Cancer |
| MDA-MB-231 | Breast | Mesenchymal (Epithelial Origin) | Carcinoma | Triple-Negative Breast Cancer (Mesenchymal subtype) |
| MCF10A | Breast | Epithelial | Non-Tumourigenic | Normal<br>Spontaneously immortalised |
| KMBC2 | Bladder | Epithelial | Carcinoma | Bladder Cancer |
| Jurkat | Blood | Lymphoblast | Leukaemia | T-cell acute lymphoblastic leukaemia (T-ALL) |
| BB3RC1 | Breast | Epithelial (PDX) | Carcinoma | ERα+ Breast Cancer |
| BB3RC29 | Breast | Epithelial (PDX) | Carcinoma | ERα+ Breast Cancer |
| HBCx34 | Breast | Epithelial (PDX) | Carcinoma | ERα+ Breast Cancer |

**Table S1** Features of normal, established cell line, and tumour models used in this study.

### Table S2

| Catalogue number | Description | Format | Vendor |
|---|---|---|---|
| 565388 | Fixable Viability Stain 780 | | BD Biosciences |
| 563423 | CD3ε | PE-Cy™7 | BD Biosciences |
| 562424 | CD4 | BV421 | BD Biosciences |
| 555482 | CD45 | FITC | BD Biosciences |
| 566852 | CD8 | APC | BD Biosciences |
| 562691 | CD14 | PE | BD Biosciences |

**Table S2** The fluorophore-conjugated antibodies and dye used for the immunotyping of PBMC and isolated CD4+ T-lymphocytes.

### Table S3

| | Breast Epithelial | CD4+ | Urothelial | Jurkat | KMBC2 | MDA-MB-231 | MCF7 |
|---|---|---|---|---|---|---|---|
| SFI1 | -0.1369 | 0.9674 | -0.3932 | 0.5563 | -0.0351 | -0.2973 | 0.0130 |
| NFIB | 3.7122 | -0.6498 | 0.1065 | -1.3556 | 0.5753 | -0.0209 | 0.5070 |
| COPE | 1.6458 | 3.6295 | 2.5420 | 1.6505 | 1.3990 | 1.4543 | 1.9106 |
| FOXO3 | 1.4517 | 0.1260 | 0.1737 | 0.2395 | 0.6369 | 0.1421 | 0.1397 |
| ZEB2 | -0.5646 | -0.0688 | -0.6363 | -1.1968 | -1.3886 | -0.5134 | -1.5999 |
| ACSL3 | 1.3597 | 0.2619 | 1.6830 | 0.8749 | 1.2587 | 0.9119 | 0.8584 |
| ACSL4 | -0.4641 | -0.0071 | -0.0405 | 0.8620 | 0.5532 | 1.2635 | -1.3697 |
| BCAR3 | 0.0623 | -0.5943 | -0.4659 | -0.9467 | 0.4747 | 1.3152 | 0.2469 |
| ESR1 | 1.1190 | -0.4936 | -0.7640 | -1.3123 | -0.6551 | -1.7627 | 1.3304 |
| GNAI2 | 0.5859 | 3.9261 | 2.2024 | 2.1964 | 1.1807 | 2.1225 | 1.7786 |
| NR3C2 | 0.1175 | 0.1706 | -0.6585 | -0.9137 | -1.3190 | -0.6092 | -1.5272 |
| NR2F6 | -0.0599 | -0.4592 | 1.7833 | 0.2630 | 1.1153 | 0.5752 | 1.1069 |
| NR2F1 | -0.6707 | -0.6526 | -0.7756 | -1.2620 | -0.0221 | 0.4818 | -1.4154 |
| SKI | -0.2324 | 0.2562 | -0.2586 | 0.6168 | 0.7925 | 1.0226 | 0.4623 |
| RARG | -0.4259 | -0.3036 | -0.1093 | -0.0991 | 1.4318 | 0.2149 | 1.3398 |
| TFEB | -0.5523 | -0.4580 | -0.6243 | -0.2913 | -0.4204 | -0.9869 | -0.9168 |
| RXRA | -0.4788 | -0.4679 | 1.9936 | -0.2682 | 1.5368 | 0.4871 | 1.1936 |
| RXRG | -0.7743 | -0.6540 | -0.7992 | -1.3949 | -1.5592 | -1.7945 | -1.5999 |
| HOXB8 | -0.7736 | -0.6536 | 0.2492 | -1.3430 | -1.3487 | -1.5761 | -1.5999 |
| HOXC8 | -0.7448 | -0.6532 | -0.7989 | -1.2915 | -1.3605 | -1.5091 | -0.8673 |
| HIVEP2 | 0.1458 | 1.6049 | 0.1175 | 0.2369 | -0.6076 | -0.4248 | -0.5888 |
| PPARG | -0.5581 | -0.6320 | 2.8215 | -1.0095 | 1.3432 | 0.9589 | -0.4200 |
| ZEB1 | -0.5990 | 1.3218 | -0.7550 | 1.1318 | -1.5334 | 0.8707 | -1.5283 |

| | | | | | | | |
|---|---|---|---|---|---|---|---|
| PBX1 | 1.9554 | -0.6243 | 0.9700 | -1.1715 | 1.3093 | 0.1545 | 1.3505 |
| PBX3 | -0.0954 | -0.1292 | -0.2814 | 0.6466 | 0.7413 | 0.2997 | 0.6646 |
| ETV5 | -0.5835 | -0.6487 | -0.7350 | 0.4243 | -1.5028 | 1.1668 | -1.1488 |
| ARHGAP25 | -0.7049 | 1.4999 | -0.7539 | 0.3909 | -1.5380 | -1.3153 | -1.5999 |
| CSNK1E | 0.9637 | -0.0524 | 1.3620 | 1.1411 | 0.6435 | 0.7943 | 0.7424 |
| NEK2 | -0.7647 | -0.6267 | -0.7769 | 1.2196 | 0.9537 | 1.0091 | 0.5343 |
| AGFG1 | 0.5109 | 0.3103 | 0.2043 | 1.0424 | 0.7027 | 0.6359 | 0.8052 |
| CRIP2 | 0.2810 | 0.5661 | 1.1942 | -0.3654 | 0.0811 | 0.0176 | 2.0467 |
| PKNOX1 | -0.2834 | -0.1662 | -0.2983 | 0.3926 | -0.0246 | -0.2086 | -0.0292 |
| PKNOX2 | -0.6601 | -0.6528 | -0.7968 | -1.3876 | -1.5105 | -1.7025 | -1.5999 |
| GSDMD | -0.0328 | 1.4211 | 0.4505 | 0.9229 | 0.9205 | 0.9811 | -1.4290 |
| IRX4 | -0.7133 | -0.6536 | -0.7987 | -1.3879 | -1.5592 | -1.8147 | -0.0995 |
| SATB1 | -0.5237 | 3.0739 | 0.3013 | 1.5369 | -1.0880 | -0.2415 | -1.5480 |
| SATB2 | -0.7262 | -0.6493 | -0.7565 | 0.1379 | -1.0868 | -0.1828 | -0.6467 |
| MECOM | -0.6530 | -0.6068 | 1.6799 | -0.8587 | 1.0329 | 0.6924 | -0.1263 |
| PRDM16 | -0.7360 | -0.6506 | -0.6771 | -1.2909 | -1.5484 | -1.7978 | -0.8919 |
| CDC20 | -0.7267 | -0.4260 | -0.7347 | 1.9204 | 1.7978 | 1.9379 | 1.5100 |
| FOXC1 | -0.3992 | -0.6531 | 0.5464 | -1.1755 | 0.5141 | -0.2659 | -0.4947 |
| CHAF1A | -0.6250 | -0.3656 | -0.5221 | 1.3722 | 0.8291 | 0.8431 | 0.6931 |
| SKP2 | -0.6981 | -0.2630 | -0.5350 | 1.2047 | 0.5675 | 0.2619 | 0.1887 |
| MAMLD1 | -0.6549 | -0.6194 | -0.7956 | -1.1858 | -1.5267 | 0.3546 | -0.9250 |
| KLF5 | 0.3971 | -0.5009 | 4.6565 | -0.5683 | 1.8153 | 0.4485 | 0.0216 |
| E2F2 | -0.7677 | -0.5818 | -0.7903 | 0.6607 | -0.3581 | -0.7586 | -0.1885 |
| E2F3 | -0.3170 | -0.1774 | -0.1904 | 0.7226 | 0.0959 | 0.4475 | 0.0319 |
| TEAD1 | 0.5430 | -0.6473 | 0.4439 | -0.3181 | 0.2499 | 1.1246 | 0.3084 |
| TEAD2 | -0.4037 | -0.6493 | -0.7932 | -1.3709 | -0.7065 | 0.0137 | 0.8099 |

| | | | | | | | |
|---|---|---|---|---|---|---|---|
| TEAD3 | -0.2943 | -0.6450 | -0.0728 | -0.7117 | 0.8513 | 0.6583 | 0.4137 |
| TEAD4 | -0.4798 | -0.6427 | -0.7303 | -0.7230 | 0.8539 | 0.9944 | 0.9292 |
| IQCC | -0.7304 | -0.6107 | -0.7395 | -0.0971 | -0.5628 | -0.9394 | -0.5707 |
| CDCA2 | -0.7365 | -0.5662 | -0.7631 | 0.9088 | 0.4182 | 0.5331 | 0.3814 |
| GRHL2 | 0.5776 | -0.6518 | -0.3271 | -1.3853 | 1.0621 | -1.7912 | 1.6709 |
| TAOK1 | 0.5137 | 0.6958 | 0.7599 | 0.9440 | 0.7138 | 0.8611 | 0.5895 |
| TTC21B | -0.3115 | -0.0657 | -0.5738 | 0.1809 | -0.4988 | -0.5791 | -0.5406 |
| MAML2 | 2.3737 | 2.8196 | 0.1660 | -1.2795 | -0.1671 | -0.3734 | -1.5666 |
| ZBTB44 | 0.1649 | 0.5716 | 1.0307 | 1.0318 | 0.5174 | 0.3598 | -0.1361 |
| KNL1 | -0.7242 | -0.5169 | -0.7303 | 1.4201 | 0.7453 | 0.8034 | 0.5770 |
| MITD1 | -0.2372 | 1.0965 | -0.0093 | 1.0271 | 0.7689 | 0.7259 | 0.3855 |
| ZNF516 | -0.3265 | -0.4663 | -0.5038 | -0.4705 | -1.5073 | -0.4712 | -0.2643 |
| TFAP2C | -0.0422 | -0.6540 | -0.2841 | 0.2713 | 0.3149 | 0.7568 | 1.5272 |
| TFAP2A | 1.0699 | -0.6528 | 0.1531 | -0.3658 | 0.7885 | -0.1356 | 0.9328 |
| TBC1D31 | -0.6806 | -0.3287 | -0.7333 | 0.7812 | 0.0206 | 0.2309 | 0.0128 |
| LCOR | 0.5267 | 0.6260 | 1.4322 | 0.6113 | -0.0781 | 0.3871 | 0.0850 |
| CCDC77 | -0.6276 | -0.3776 | -0.6854 | 0.8726 | 0.2286 | 0.2687 | -0.0885 |
| TRIM9 | -0.7296 | -0.6456 | -0.7340 | -0.4362 | -0.8410 | -1.4759 | -1.5584 |
| WWTR1 | 1.1435 | -0.6033 | -0.1287 | -1.3228 | 0.6999 | 1.0145 | -0.0260 |
| BCL11A | -0.3153 | -0.5950 | -0.1681 | 0.5456 | -1.5508 | -0.9791 | -1.5999 |
| UPF2 | 0.7751 | 1.7222 | 0.5662 | 0.6304 | 0.3208 | 0.2160 | 0.6247 |
| CPAP | -0.6361 | -0.4639 | -0.6336 | 0.5750 | 0.0534 | -0.1191 | 0.0417 |
| DCAF16 | -0.2985 | 0.2583 | -0.1918 | 0.6924 | 0.3634 | 0.5107 | -0.2159 |
| GTSE1 | -0.7354 | -0.5559 | -0.7674 | 1.3258 | 0.7650 | 1.0016 | 0.5380 |
| GRHL1 | 0.2772 | -0.6407 | 0.6181 | -0.4618 | 0.3636 | -1.6963 | 0.6003 |
| POLK | 0.0205 | 0.3014 | -0.1803 | 0.5386 | -0.1798 | -0.1145 | -0.4174 |

| | | | | | | | |
|---|---|---|---|---|---|---|---|
| TRPS1 | 5.2738 | 0.0419 | -0.7333 | -1.3333 | -1.5592 | -0.5264 | 1.0464 |
| IKZF3 | -0.6774 | 1.6775 | -0.5328 | -0.1803 | -1.0961 | -1.7757 | -1.5725 |
| ZMIZ1 | -0.0050 | -0.2657 | 0.1894 | 0.5846 | -0.0416 | 0.7469 | 0.5354 |
| SOX13 | -0.5216 | -0.6249 | -0.3845 | -1.1445 | 0.5461 | 0.5419 | 0.5649 |
| RNF114 | 0.8665 | 1.4332 | 0.4134 | 1.3095 | 1.0134 | 1.1934 | 1.8072 |
| AR | 0.5499 | -0.6456 | -0.2346 | -1.3949 | -1.5554 | -0.0242 | 0.2412 |
| PGR | 0.0336 | -0.6529 | -0.7981 | -1.3949 | -1.5592 | -1.7722 | 0.2118 |
| HOXA5 | -0.6070 | -0.6517 | -0.4398 | 0.2545 | 0.2947 | -1.8147 | -0.2461 |
| FOXP3 | -0.7587 | -0.2723 | -0.7934 | -1.0522 | -1.3821 | -1.7359 | -1.2463 |
| SMARCD3 | -0.4370 | -0.4920 | -0.6225 | 0.6499 | -0.9777 | -0.7017 | -0.1012 |

**Table S3** z-score normalised transcript abundance for proteins most strongly associated with tissue-specific GR interactions.

### Table S4

| **Identifier** | **Antibody Target** |
|---|---|
| SRX1161165 | NR3C1 |
| SRX1161166 | NR3C1 |
| SRX1161167 | NR3C1 |
| SRX19024866 | NR3C1 |
| SRX19024868 | NR3C1 |
| SRX19024870 | NR3C1 |
| SRX19024878 | NR3C1 |
| SRX19024879 | NR3C1 |
| SRX19024880 | NR3C1 |
| SRX19024868 | NR3C1 |
| SRX19024870 | NR3C1 |
| SRX19024878 | NR3C1 |
| SRX19024879 | NR3C1 |
| SRX19024880 | NR3C1 |
| SRX1012654 | PGR |
| SRX1012653 | PGR |
| SRX1012655 | PGR |
| SRX10365514 | AR |
| SRX10365515 | AR |
| SRX19024872 | ESR1 |
| SRX1012630 | ESR1 |
| SRX19024874 | ESR1 |
| SRX4451215 | ESR1 |
| SRX1012629 | ESR1 |
| SRX10365532 | ESR1 |
| SRX4451217 | ESR1 |
| SRX1531796 | ESR1 |
| SRX4451216 | ESR1 |
| SRX1161171 | ESR1 |
| SRX3080213 | ESR1 |
| SRX3583203 | ESR1 |
| SRX20282946 | ESR1 |
| SRX3110718 | ESR1 |
| SRX3080212 | ESR1 |
| SRX5287706 | ESR1 |
| SRX2922226 | ESR1 |
| SRX1012694 | PGR |
| SRX1012683 | ESR1 |
| SRX1012682 | ESR1 |

**Table S4** Comprehensive list of SRX identifiers for all ChIP-seq samples used in the ChIP-Atlas colocalisation analysis of GR (NR3C1) binding with AR, ERα and PGR in MCF7 cells.

Table S5

| Identifier | Antibody Target |
|---|---|
| SRX3070467 | NR3C1 |
| SRX3070469 | NR3C1 |
| SRX3070471 | NR3C1 |
| SRX1012659 | PGR |
| SRX1012658 | PGR |
| SRX1012660 | PGR |
| SRX1012654 | PGR |
| SRX1012653 | PGR |
| SRX1012655 | PGR |
| SRX10365514 | AR |
| SRX10365515 | AR |
| SRX10365534 | ESR1 |
| SRX10365557 | ESR1 |
| SRX10365533 | ESR1 |
| SRX1012632 | ESR1 |
| SRX1585243 | ESR1 |
| SRX1161168 | ESR1 |
| SRX7846778 | ESR1 |
| SRX5465793 | ESR1 |
| SRX19024872 | ESR1 |
| SRX1012630 | ESR1 |
| SRX19024874 | ESR1 |
| SRX2943589 | ESR1 |
| SRX2943585 | ESR1 |
| SRX4451215 | ESR1 |
| SRX10365536 | ESR1 |
| SRX1012633 | ESR1 |
| SRX1012629 | ESR1 |
| SRX10365532 | ESR1 |
| SRX3110722 | ESR1 |
| SRX371468 | ESR1 |
| SRX3080214 | ESR1 |
| SRX1012631 | ESR1 |
| SRX3123829 | ESR1 |
| SRX2943591 | ESR1 |
| SRX194572 | ESR1 |
| SRX4451217 | ESR1 |
| SRX371469 | ESR1 |
| SRX19024875 | ESR1 |
| SRX1585242 | ESR1 |
| SRX5323878 | ESR1 |

| | |
|---|---|
| SRX1012628 | ESR1 |
| SRX3229134 | AR |

**Table S5** Comprehensive list of SRX identifiers for all ChIP-seq samples used in the ChIP-Atlas colocalisation analysis of GR (NR3C1) binding with AR, ERα and PGR in MCF10A cells.

### Table S6

| **Identifier** | **Antibody Target** |
|---|---|
| SRX3229187 | GR |
| SRX3229190 | PGR |
| SRX3229161 | ESR1 |
| SRX186804 | ESR1 |
| SRX3229172 | AR |
| SRX3229168 | AR |
| SRX3229173 | AR |
| SRX5702625 | ESR1 |
| SRX5702624 | ESR1 |
| SRX5493769 | ESR1 |
| SRX5493763 | ESR1 |
| SRX5323930 | ESR1 |
| SRX5323929 | ESR1 |
| SRX4116034 | ESR1 |
| SRX3229165 | ESR1 |
| SRX3229164 | ESR1 |
| SRX3229163 | ESR1 |
| SRX3229156 | ESR1 |
| SRX3229155 | ESR1 |
| SRX3229154 | ESR1 |
| SRX3229153 | ESR1 |
| SRX3229152 | ESR1 |
| SRX3229151 | ESR1 |
| SRX3229150 | ESR1 |
| SRX3229149 | ESR1 |
| SRX3229148 | ESR1 |
| SRX3229147 | ESR1 |
| SRX3229146 | ESR1 |
| SRX3229144 | ESR1 |
| SRX3229143 | ESR1 |
| SRX3229142 | ESR1 |
| SRX3229140 | ESR1 |
| SRX3229139 | ESR1 |
| SRX3229136 | ESR1 |
| SRX3229132 | ESR1 |
| SRX3229121 | ESR1 |
| SRX2897201 | ESR1 |
| SRX186824 | ESR1 |
| SRX186821 | ESR1 |
| SRX186813 | ESR1 |
| SRX186810 | ESR1 |
| SRX186807 | ESR1 |

| | |
|---|---|
| SRX3229157 | ESR1 |
| SRX3229145 | ESR1 |
| SRX2897197 | ESR1 |

**Table S6** Comprehensive list of SRX identifiers for all ChIP-seq samples used in the ChIP-Atlas colocalisation analysis of GR (NR3C1) binding with AR, ERα and PGR in Tumour and PDX samples.

### Table S7

| Identifier | Antibody Target |
|---|---|
| SRX1161165 | NR3C1 |
| SRX1161166 | NR3C1 |
| SRX1161167 | NR3C1 |
| SRX19024866 | NR3C1 |
| SRX19024868 | NR3C1 |
| SRX19024878 | NR3C1 |
| SRX19024879 | NR3C1 |
| SRX19024880 | NR3C1 |
| SRX13298925 | ZEB1 |
| SRX13298923 | ZEB1 |
| SRX13298924 | ZEB1 |

**Table S7** Comprehensive list of SRX identifiers for all ChIP-seq samples used in the ChIP-Atlas colocalisation analysis of GR (NR3C1) binding with ZEB1 in MCF7 cells.

### Table S8

| Identifier | Antibody Target |
|---|---|
| SRX1161165 | NR3C1 |
| SRX1161166 | NR3C1 |
| SRX1161167 | NR3C1 |
| SRX19024866 | NR3C1 |
| SRX19024868 | NR3C1 |
| SRX19024878 | NR3C1 |
| SRX19024879 | NR3C1 |
| SRX19024880 | NR3C1 |
| SRX3401273 | TRPS1 |
| SRX6098144 | TRPS1 |
| SRX190316 | TEAD4 |

**Table S8** Comprehensive list of SRX identifiers for all ChIP-seq samples used in the ChIP-Atlas colocalisation analysis of GR (NR3C1) binding with TRPS and TEAD4 in MCF7 cells.

### Supplementary Methods

#### Data-dependent acquisition mass spectrometry analysis (Orbitrap Fusion)

Peptides were loaded onto an mClass nanoflow UPLC system (Waters) equipped with a nanoEaze M/Z Symmetry 100 Å $C_{18}$, 5 µm trap column (180 µm x 20 mm, Waters) and a PepMap, 2 µm, 100 Å, $C_{18}$ EasyNano nanocapillary column (75 mm x 500 mm, Thermo). The trap wash solvent was aqueous 0.05% (v:v) trifluoroacetic acid and the trapping flow rate was 15 µL/min. The trap was washed for 5 minutes before switching flow to the capillary column. Separation used gradient elution of two solvents: solvent A, aqueous 0.1% (v:v) formic acid; solvent B, acetonitrile containing 0.1% (v:v) formic acid. The flow rate for the capillary column was 330 nL/min and the column temperature was 40°C. The linear multi-step gradient profile was: 3-10% B over 7 minutes, 10-35% B over 30 minutes, 35-99% B over 5 minutes and then proceeded to wash with 99% solvent B for 4 minutes. The column was returned to initial conditions and re-equilibrated for 15 minutes before subsequent injections.

The nanoLC system was interfaced with an Orbitrap Fusion Tribrid mass spectrometer (Thermo) with an EasyNano ionisation source (Thermo). Positive ESI-MS and $MS^2$ spectra were acquired using Xcalibur software (version 4.0, Thermo). Instrument source settings were: ion spray voltage, 1,900 V; sweep gas, 0 Arb; ion transfer tube temperature; 275°C. $MS^1$ spectra were acquired in the Orbitrap with: 120,000 resolution, scan range: *m/z* 375-1,500; AGC target, $4e^5$; max fill time, 100 ms. Data dependent acquisition was performed in top speed mode using a 1 s cycle, selecting the most intense precursors with charge states > 1.  Easy-IC was used for internal calibration. Dynamic exclusion was performed for 50s post precursor selection, and a minimum threshold for fragmentation was set at $5e^3$. $MS^2$ spectra were acquired in the linear ion trap with: scan rate, turbo; quadrupole isolation, 1.6 *m/z*; activation type, HCD; activation energy: 32%; AGC target, $5e^3$; first mass, 110 *m/z*; max fill time, 100 ms. Acquisitions were arranged by Xcalibur to inject ions for all available parallelisable time.

#### Data-dependent acquisition mass spectrometry analysis (timsTOF HT)

Peptides were loaded onto EvoTip Pure tips for desalting and as a disposable trap column for nanoUPLC using an EvoSep One system. A pre-set EvoSep 100 SPD gradient (from Evosep One HyStar Driver 2.3.57.0) was used with an 8 cm EvoSep $C_{18}$ Performance column (8 cm x 150 mm x 1.5 mm).

The nanoUPLC system was interfaced to a timsTOF HT mass spectrometer (Bruker) with a CaptiveSpray ionisation source (Source). Positive PASEF-DDA, ESI-MS and $MS^2$ spectra were acquired using Compass HyStar software (version 6.2, Bruker). Instrument source settings were: capillary voltage, 1,500 V; dry gas, 3 l/min; dry temperature; 180°C. Spectra were acquired between *m/z* 100-1,700. TIMS settings were: 1/K0 0.6-1.60 V.s/cm$^2$; Ramp time, 100 ms; Ramp rate 9.42 Hz. Data-dependent acquisition was performed with 10 PASEF ramps and a total cycle time of 1.17 s. An intensity threshold of 2,500 and a target intensity of 20,000 were set with active exclusion applied for 0.4 min post precursor selection. Collision energy was interpolated between 20 eV at 0.6 V.s/cm$^2$ to 59 eV at 1.6 V.s/cm$^2$.

#### DDA Database Searching

Data were searched using FragPipe (v23.0) against the human subset of SwissProt appended with common proteomic contaminants. Searches were set to require specific trypsin specificity. Set tolerances allowed for fragment ion mass precision of 0.50 Da (Orbitrap Fusion) or 20 ppm (Bruker timsTOF HT) and a parent ion tolerance of 3.0 ppm (Orbitrap Fusion) or 10 ppm (Bruker timsTOF HT). Searches were performed with a target of 1% FDR. Non-normalised peak intensities were extracted using IonQuant specifying FDR < 0.01 and a 3 ppm (Orbitrap Fusion) or 10 ppm (Bruker timsTOF HT) tolerance for match between runs. Protein identifications were filtered to require a minimum of two peptides.
